# Ancient introgression explains mitochondrial genome capture and mitonuclear discordance among South American collared *Tropidurus* lizards

**DOI:** 10.1101/2025.04.25.650633

**Authors:** Matheus M. A. Salles, André L. G. Carvalho, Adam D. Leaché, Nicolas Martinez, Frederick Bauer, Martha Motte, Viviana Espínola, Miguel T. Rodrigues, Carla Piantoni, Marcio R. Pie, André Olivotto, Guarino R. Colli, Erik L. Choueri, Fernanda P. Werneck, Fabricius M. C. B. Domingos

## Abstract

Mitonuclear discordance—evolutionary discrepancies between mitochondrial and nuclear DNA phylogenies—can arise from various factors, including introgression, incomplete lineage sorting, recent or ancient demographic fluctuations, sex-biased dispersal asymmetries, among others. Understanding this phenomenon is crucial for accurately reconstructing evolutionary histories, as failing to account for discordance can lead to misinterpretations of species boundaries, phylogenetic relationships, and historical biogeographic patterns. We investigate the evolutionary drivers of mitonuclear discordance in the *Tropidurus spinulosus* species group, which contains nine species of lizards inhabiting open tropical and subtropical environments in South America. Using a combination of population genetic and phylogenomic approaches applied to mitochondrial and nuclear data, we identified different instances of gene flow that occurred in ancestral lineages of extant species. Our results point to a complex evolutionary history marked by prolonged isolation between species, demographic fluctuations, and potential episodes of secondary contact with genetic admixture. These conditions likely facilitated mitochondrial genome capture while diluting signals of nuclear introgression. Furthermore, we found no strong evidence supporting incomplete lineage sorting or natural selection as primary drivers of the observed mitonuclear discordance. Therefore, the unveiled patterns are most consistent with neutral demographic processes, coupled with ancient mitochondrial introgression, as the main factors underlying the mismatch between nuclear and mitochondrial phylogenies in this system. Future research could further explore the role of other demographic processes, such as asymmetric sex-biased dispersal, in shaping these complex evolutionary patterns.

## Introduction

Mitochondrial and nuclear genomes evolve independently, through distinct processes, often leading to conflicting genetic patterns—a phenomenon known as mitonuclear discordance. Cases of mitonuclear discordance are increasingly reported across a wide range of animal species. Recent examples can be found among mammals (Good et al. 2015; Phuong et al. 2016; Seixas et al. 2018; Fedorov et al. 2022), birds (Andersen et al. 2021; DeRaad et al. 2023), reptiles (Firneno et al. 2021), amphibians (Zieliński et al. 2013; Dufresnes et al. 2020; Rancilhac et al. 2021), and insects (Hinojosa et al. 2019; Dong et al., 2023). Generally, conflicting patterns between nuDNA and mtDNA arise from processes such as introgression, demographic fluctuations, sex-biased dispersal asymmetries, incomplete lineage sorting (ILS), human introductions, or *Wolbachia* infection in insects (see reviews by Toews and Brelsford 2012; Bonnet et al. 2017; Deprés 2019). Determining whether mitonuclear discordance is driven by any of these factors requires a clear understanding of the genealogies among different loci, the demographic history of populations, and their historical introgression/migration patterns and rates. Notably, these complexities often present significant empirical challenges in resolving the underlying causes of discordance.

Most cases of mitonuclear discordance are attributed to mtDNA introgression (Toews and Brelsford 2012), although the extent of introgression varies widely among animal groups. For instance, Zozaya et al. (2024) showed that, despite frequent hybrid formation across multiple contact zones in Australian *Heteronotia* geckos, mtDNA exchange can be limited to narrow geographic areas, indicating that hybridization does not necessarily result in widespread mitochondrial replacement. When discordance is attributable to introgression, it is most often linked to processes such as gene flow across hybrid zones (Zieliński et al. 2013; Good et al. 2015; Dufresnes et al. 2020; Mao and Rossiter 2020) or mitochondrial capture (Andersen et al. 2021). The latter refers specifically to a form of complete introgression, where the mitochondrial genome of one species is entirely replaced by another’s (Zieliński et al. 2013). Mitochondrial capture can occur at various stages of the speciation process and between distantly related species (Rancilhac et al. 2021). Additionally, mitochondrial introgression can take place with minimal or no accompanying nuDNA introgression (Zieliński et al. 2013; Good et al. 2015; Andersen et al. 2021; Rancilhac et al. 2021; Fedorov et al. 2022), making it challenging to detect introgressed nuclear and mitochondrial markers simultaneously (Bonnet et al. 2017; Seixas et al. 2018). Therefore, ILS is another phenomenon that cannot be overlooked in the discussion of mitonuclear discordance, given the challenges in distinguishing it from introgression (Andersen et al. 2021; DeRaad et al. 2023). While some studies have identified ILS as a primary driver of mitonuclear discordance, these remain relatively rare (e.g., Firneno et al. 2021).

Demographic factors, such as population size fluctuations, particularly at the periphery of a species’ geographic range, can also contribute to mitonuclear discordance (Phuong et al. 2016; Dufresnes et al. 2020; Fedorov et al. 2022). In this scenario, neutral processes such as genetic drift can help in the fixation of mtDNA haplotypes at range edges of expanding populations (Bonnet et al. 2017). Even in situations where introgression involves both mtDNA and nuclear loci, mtDNA is expected to reach fixation faster, since it has a smaller effective population size and is non-recombining (Moore 1995). Following demographic expansions and secondary contact, nuclear genomes can fully recombine, thereby diluting historical introgression signals, whereas divergent, non-recombining mitochondrial haplotypes may persist.

Finally, selection can also play a role in generating mitonuclear discordance (Toews and Brelsford 2012). If a beneficial mitochondrial mutation arises but is initially incompatible with the nuclear genetic background of a population, selection may favor nuclear variants that restore mitonuclear compatibility. This compensatory selection could lead to a scenario where the advantageous mitochondrial mutation spreads widely, while the nuclear DNA follows a different evolutionary trajectory. Empirical evidence robustly demonstrates that natural selection can drive mtDNA introgression between populations through hybridization, even in the absence of corresponding nuDNA introgression, provided there is even a minor selective advantage (Excoffier et al. 2009; Zieliński et al. 2013; Good et al. 2015; Phuong et al. 2016). As a result, the mitochondrial genome with the highest fitness may introgress into another species, regardless of whether it originates from the resident or the invading population.

Here, we investigate the drivers of mitonuclear discordance in a biological system with known occurrences of this phenomenon, the lizard family Tropiduridae (e.g., *Microlophus*: Benavides et al. 2007; 2009; *Tropidurus*: Carvalho et al. 2016). Specifically, we explored the evolutionary history of nine lineages within the *T. spinulosus* group using a multi-locus genomic approach. This diverse group of lizards is distributed across open-vegetation tropical and subtropical environments in South America and predominantly inhabit rocky and arboreal habitats (Frost et al. 1998; Carvalho 2013, 2016; Carvalho et al. 2013). Although the taxonomy of the *T. spinulosus* group has been relatively stable, new species continue to be described (e.g., Carvalho 2016). Additionally, our research includes all taxa presently admitted to the group plus one undescribed species, which is currently undergoing formal description (Carvalho et al. in prep) and is referred to as *Tropidurus* sp. nov. in this study.

We integrated mitochondrial genomes, nuclear ultraconserved elements (UCEs), and single-nucleotide polymorphisms (SNPs) derived from UCEs to assess whether the mitochondrial genealogy corresponded to reciprocally monophyletic nuclear lineages. To determine the potential sources of the mitonuclear discordance, we examined the relative contributions of neutral and selective processes. Specifically, we first compared mtDNA and nuclear phylogenies to identify conflicts and potential introgression patterns, using methods that either account for ILS or do not. In a scenario where these approaches yield largely congruent results, ILS would not play a significant role in the observed mitonuclear discordance. Second, we evaluated whether mtDNA introgression was accompanied by detectable levels of nuDNA introgression, as most cases of discordance are attributed to processes such as gene flow across hybrid zones. We also assessed signals of positive selection on mtDNA to evaluate potential adaptive processes in shaping mitonuclear discordance. Finally, as demographic processes can contribute to emerging phylogenetic discordant patterns as well, we examined historical changes in population size to assess their role in shaping the observed discordance, as this would align with expectations under a non-adaptive scenario.

## Methods

### Sampling and Lab work

The phylogenomic approaches used in this study were based on anchored sequencing methods, specifically Ultraconserved Elements (UCE) as described by Faircloth et al. (2012). We sequenced a total of 43 individuals (Supplementary Material, Table S9), representing all species of the *Tropidurus spinulosus* group. RAPiD Genomics LLC (Gainesville, FL) was tasked with sample library preparation and target enrichment of UCEs using the tetrapod 5k probe set (Faircloth et al. 2012), followed by multiplexed paired-end (PE) sequencing (2 × 100 bp) of UCEs on an Illumina HiSeq 3000 PE100 platform. Our samples are mainly distributed around the Pantanal Basin, with some extending to dry environments of Paraguay and Argentina (Figure 1).

**Figure 1.**
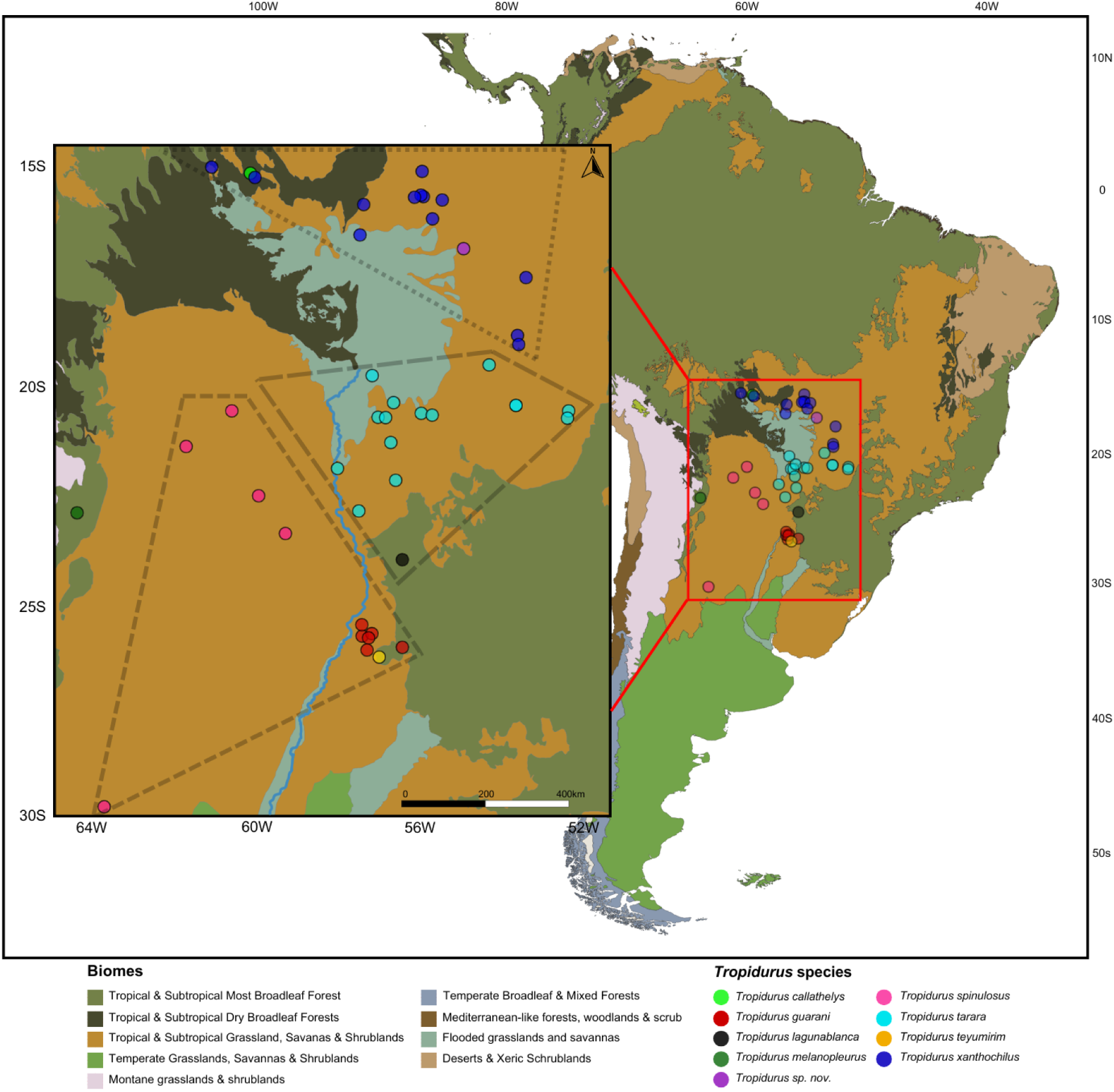
Sample distribution with species assignments based on our concatenated phylogenetic analysis. The map at a larger scale shows the distribution of samples across various South American ecoregions, according to the classification of <u>“Terrestrial Ecoregions of the World” adopted by the World Wildlife Fund</u>. For a detailed review of the resolution and delimitation differences in these categories, see Olson et al. (2001). The smaller-scale map highlights the specific region where the analyzed samples are distributed. Phylogenetically identified groups are marked with three distinct dashed line patterns, emphasizing their allopatric distribution: (1) the *T. spinulosus* + *T. guarani* + *T. teyumirim clade* (multi-species), (2) the *T. tarara clade* (with *T. lagunablanca* tentatively considered a synonym of *T. tarara*), and (3) the *T. xanthochilus* + *T. sp. nov.* clade. The map in the smaller box also shows part of the course of the Paraguay River (light blue), which separates the *T. spinulosus* lineages to the west from *T. guarani* and *T. tarara* to the east.

### UCEs and mtDNA pipeline

We assembled reads using the Phyluce v1.7.1 pipeline (Faircloth 2016). Demultiplexed reads were cleaned to remove low-quality bases and adapter sequences in Trimmomatic (Bolger et al. 2014), using the wrapper program illumiprocessor in Phyluce. To accelerate assembly and render challenging datasets tractable, we normalized read depth to a minimum of 5 with the bbnorm.sh script from the BBTools suite (Bushnell 2018). Trimmed and normalized reads were inspected for quality and adapter contamination using FastQC v0.11.9 (Andrews 2010), and then assembled into contigs with SPAdes v3.15.5 (Bankevich et al. 2012). We used Phyluce to align assembled contigs back to their associated UCE loci, remove duplicate matches, create a taxon-specific database of contig-to-UCE matches, and extract UCE loci for all individuals. Specifically, contigs matching UCE loci were identified and extracted using the program LastZ v1.0 (Harris 2007) within Phyluce, both in incomplete and complete (75% and 95% of completeness) matrices (see Faircloth 2016). After probes and UCEs were matched, we aligned UCE contigs with MAFFT v7.471 (Katoh and Standley 2013) using specific customized settings (-globalpair, --maxiterate 1000, --adjustdirection), and trimmed the resulting alignments using the internal-trimming algorithm (Gblocks: Castresana 2000). As a final step, AMAS (Borowiec 2016) was used to compute final summary statistics for all alignments.

Due to the high abundance of mtDNA in samples and the less-than-perfect efficiency of target enrichment methods, mitochondrial data (including entire mitogenomes) are often generated as a byproduct of the UCE sequencing process (Allio et al. 2020). Thus, we extracted mtDNA from our contig assembly produced through Phyluce using MitoFinder (Allio et al. 2020), which can find and extract multiple mitochondrial genes or even entire mitogenomes from assembled sets of bulk sequences. In general terms, mitochondrial sequences were aligned following the same procedure applied to UCE data, as described above.

### SNP calling workflow

We selected the sample with the largest number of UCE loci among all individuals (‘MTR 29586’, with 4800 loci) retrieved during the extraction step described above as the primary reference during the process of calling SNPs as recommended in the Zarza et. al. (2016) pipeline to extract SNPs from UCE reads. Then, we followed the workflow developed by Harvey et al. (2016) to sequence capture data from population-level samples. The first step involves BWA, which we used to map raw reads from individuals to contigs (Li et al. 2009; Li 2013). Thus, SAM files were converted to BAM with SAMtools (Li et al. 2009). Alignments were checked for BAM format violations, read group header information was added, and PCR duplicates were marked for each individual using [Picard (v. 1.106)](https://broadinstitute.github.io/picard/). The resulting BAM files for each individual in a lineage were merged into a single file with Picard, which was then indexed with SAMtools. The Genome Analysis Toolkit (GATK; v. 3.4-0; McKenna et al. 2010) was used to locate and realign around indels, which was followed by calling SNPs using the ‘UnifiedGenotyper’ tool in GATK. SNPs and indels were then annotated, and indels were masked. Finally, we used GATK to restrict our datasets to high-quality SNPs (Q30 filter) and performed read-backed phasing. After that, we used the R packages SNPfiltR (DeRaad 2022) and vcfR (Knaus and Grunwald 2017) to visually and iteratively filter our SNP dataset based on specific quality and completeness metrics. For analyses requiring unlinked SNPs, a linkage disequilibrium-based filtering approach was applied using PLINK (Purcell et al. 2007) with fixed parameters (--indep-pairwise 50 10 0.5). At the end of this process, two datasets were generated. The first, referred to as the ‘*unlinked SNP dataset I*’, included 42 samples and excluded the *Tropidurus melanopleurus* sample, retaining only one sample external to the largest clade within the group, namely *T. callathelys* (used in D-suite analysis; 40,265 SNPs, 25.34% missing data). The second one, ‘*unlinked SNP dataset II’,* with 37 samples, excluding the *T. melanopleurus, T. callathelys, T. teyumirim*, and *Tropidurus* sp. nov. samples, and assigning the *T. lagunablanca* sample as *T. tarara* (used in Stairway Plot analysis; 27,454 SNPs, 24.71 % missing data).

### Gene and species tree analysis

Gene trees for all UCE loci and the mtDNA locus were inferred using IQ-TREE v2.2.6 (Minh et al. 2020), and support was inferred using 1000 ultrafast bootstraps (≥ 95 considered as strong support). For partitioned ML analyses, ModelFinder (Kalyaanamoorthy et al. 2017), part of the IQ-TREE package, was used to select the best-fit model for each partition, followed by tree reconstruction (-m TESTNEWMERGE option), allowing partitions to have different evolutionary rates (-spp option). Support was inferred using 1000 ultrafast bootstraps. Additionally, species trees were inferred for the UCE dataset using ASTRAL v5.15.4 (Zhang et al. 2018), which does this from gene trees and provides internal branch lengths in coalescent units of gene tree discordance, as well as branch support values in the form of local posterior probabilities (LPPs). In our ASTRAL analyses, we used the topology resulting from the partitioned approach produced through IQ-TREE as a reference. Finally, we used BPP v.4.6 (Flouri et al., 2018) to estimate a species tree for the *Tropidurus spinulosus* group, based on the full dataset of 820 UCE loci.

In the case of mitochondrial data, besides partitioning by gene, the alignment was also partitioned by third codon positions (in the case of coding sequences), and we did an additional analysis including two outgroup species to evaluate the monophyly of the *Tropidurus spinulosus* group — *Plica plica* and *T. torquatus* (Figure S6; GenBank accession numbers: AB218961 for *P. plica*, and KU245273, KU245300, KU245090 and KU245062 for *T. torquatus*). We present only the trees (both based on nuDNA and mtDNA) featuring species from the *T. spinulosus* group (our in-group) in the results section. In all cases, trees were manually rooted in *T. melanopleurus* using FigTree (Rambaut 2014), following the topology obtained in the previously described analysis.

### Divergence time estimation

A time tree was inferred using two different approaches. The first was the RelTime-Branch Lengths method (Tamura et al., 2018) implemented in MEGA11 (Tamura et al., 2021), which infers divergence times from a previously estimated phylogenetic tree with branch lengths. In this case, the time-tree was computed using the UCE-based phylogenetic tree (complete matrix) with one calibration constraint, namely the node separating *Tropidurus callathelys* from all the other species in the group (and using *T. melanopleurus* as the outgroup). We used 10.74 Ma as the reference value for this constraint, following the estimates obtained by Zheng and Wiens (2016) regarding the node separating *T. callathelys* and the rest of the species in our ingroup. The method described in Tao et al. (2020) was used to estimate confidence intervals and set a normal distribution (mean = 10.74 and standard deviation = 0.5) on the node for which calibration densities were provided.

Alternatively, divergence time estimates were also estimated using MCMCTree v4.10 (dos Reis and Yang 2019), which uses an approximate likelihood approach. In this case, the 100 most clocklike UCE loci within our dataset were selected, particularly the loci with the least root-to-tip variance. Clock-likeness was assessed using SortaDate (Smith et al. 2018), which measures root-to-tip variance within gene trees and then sorts them from highest to lowest variance. The calibration in this case was performed by constraining the age of the same node described above to be between 10 and 11 Ma, using a fixed topology. Molecular clock estimates were obtained using the independent rates model, and we used the HKY85 model. The gamma prior on the mean substitution rate for partitions (rgene_gamma) was set to G (7, 9.75), which means a substitution rate of approximately 0.00717 substitutions/site/Ma (value available for the phylogenetically closest species to *Tropidurus* in the germline mutation rate study of Bergeron et al. 2023). The gamma rate variance (sigma2_gamma) was specified with G (1, 10). Two independent runs were performed, each consisting of 550.000 MCMC iterations, with 10.000 as burn-in, ‘nsample’ = 100.000, and ‘sampfreq’ = 5. The ESS of the MCMCTree runs were examined in Tracer v.1.7 (Rambaut et al. 2018) to determine convergence, and values > 200 were retained.

### Gene flow and phylogenetic network estimation

Failing to consider the potential influence of gene flow and introgressive hybridization among species can negatively impact phylogenetic inference (e.g., Solís-Lemus and Ané 2016; Solís-Lemus et al. 2016). Therefore, given the mitonuclear discordance observed in the *Tropidurus spinulosus* clade, we evaluated the potential for reticulate evolution in our dataset using five different approaches: (i) the phylogenetic networks applying quartets (SNaQ) method implemented in PhyloNetworks (Solís-Lemus and Ané 2016; Solís-Lemus et al. 2017); (ii) the maximum pseudolikelihood estimate (MPL) method implemented in PhyloNet (Wen et al. 2018); (iii) the extension of the typical tree-based model to general networks, assuming no ILS and independent loci, available in the NetRAX program (Lutteropp et al. 2022); (iv) ABBA-BABA tests for detecting gene flow between pairs of species, through the D-suite program (Malinsky et al., 2021); and (v) the MSC-M model implemented in BPP v.4.6 (Flouri et al. 2018; 2023) to estimate migration rate between pair of species that showed signals of mitonuclear discordance. The first four methods serve as exploratory tools to identify potential reticulations or gene flow events, while the BPP-based approach directly quantifies migration rates. Regarding the phylogenetic network and the ABBA-BABA approaches, please refer to the Supplementary Material, section “*Details on phylogenetic networks methodology and results*”, for a detailed explanation of all reticulation tests.

To directly assess how gene flow differentially impacted nuclear and mitochondrial genomes, we used the migration model (MSC-M) available in BPP v.4.6 (Flouri et al. 2018; 2023). This allowed us to estimate the number of migrants per generation and the direction of gene flow for species showing the strongest mitonuclear discordance in our phylogenetic trees. This was done in two stages, either using the nuclear species tree or the mitochondrial genealogy as the guide tree. In the first case, we used only a portion of the nuDNA dataset— specifically, the 100 most clocklike UCEs selected using SortaDate (Smith et al. 2018). A second migration analysis was performed using that same dataset, but this time incorporating mitochondrial data alongside nuclear data. In other words, two analyses were conducted using the nuclear topology as the guide tree: one using nuclear markers only (100 UCE loci, 93,246 bp) and another combining nuclear data with mitochondrial data (100 UCE loci, 93,246 bp + 1 mitochondrial partition, 13,578 bp).

In both cases, we tested bidirectional migration scenarios with the following species: *Tropidurus xantochilus* ↔ *T. guarani*, *T. xanthochilus* ↔ *T. spinulosus,* and *T. xanthochilus* ↔ *T. tarara*. Scenarios involving ancestral populations or species of the following groups were also tested. For analyses using the mitochondrial genealogy as the guide tree, these included: (A) *T. tarara* and one *T. xanthochilus* group; (B) *T. guarani* and *T. spinulosus*; (C) *T. guarani, T. spinulosus*, and another *T. xanthochilus* group. When using the nuclear topology as the guide tree, the same scenarios with current species mentioned above were tested, while also considering the following scenarios involving ancestral nodes: (B) *T. guarani* and *T. spinulosus*; (D) *Tropidurus* sp. nov. and *T. xanthochilus*; (E) *T. tarara, T. guarani*, *T. spinulosus,* and *T. teyumirim*; (F) *T. guarani*, *T. spinulosus*, and *T. teyumirim.* Population sizes (θ) were also estimated and recorded for each species and ancestral node involved in the migration pairs. In all cases, we used an inverse gamma prior of G(3, 0.04) for the Θ parameter and G(3, 0.2) for the τ parameter, fixing the species model prior on the topologies described above. The gamma migration prior used was 0.1 (α = 1, β = 10). We ran analyses for 4 x 10^5^ MCMC generations, taking samples every five and using 5 x 10^3^ burn-in generations. To check for consistency of results, we conducted two independent runs for each analysis.

Furthermore, we also used POPART v. 1.7 (Leigh et al. 2015) to generate a TCS haplotype network (Clement et al. 2002), based on three mitochondrial genes - cytochrome c oxidase I, II, and III (COX1, COX2, and COX3).

### Tests for positive selection and demographic changes through time

To detect significant deviations from the hypothesis of neutral evolution in the mtDNA dataset, we tested for the evidence of positive selection using MEME (Mixed Effects Model of Evolution; Murrell et al. 2012), BUSTED (Branch-Site Unrestricted Statistical Test for Episodic Diversification; Murrell et al. 2015), aBSREL (adaptive Branch-Site Random Effects Likelihood; Smith et al. 2015), and FUBAR (Fast, Unconstrained, Bayesian Approximation; Murrell et al. 2013). MEME is designed to identify sites underlying episodic selection, BUSTED provides a gene-wide (not site-specific) test for positive selection by asking whether a gene has experienced positive selection at least one site on at least one branch, aBSREL focuses on tree branches and can be used to test if positive selection has occurred on a proportion of branches, while FUBAR uses a Bayesian approach to infer nonsynonymous (dN) and synonymous (dS) substitution rates on a per-site basis for a given coding alignment and corresponding phylogeny. All of these tests were performed utilizing the HYPHY package through the [DataMonkey server](https://datamonkey.org/), using default parameters, and the threshold of *p* < 0.05 for all four analyses. The same alignment of mitochondrial genes used for the phylogenetic analyses was utilized here, with the stop codons of each coding gene removed manually to perform the selection analyses. Also, Tajima’s D was calculated to evaluate evidence of natural selection and population size changes in the mitogenome using the R function ‘tajima.test’, from the ‘pegas’ package (Paradis 2010). Mitochondrial nucleotide diversity was estimated for populations of *T. guarani*, *T. spinulosus*, *T. tarara*, and *T. xanthochilus* using the ‘nuc.div’ function, also from the ‘pegas’ package.

Because selection tests indicated neutrality in the mtDNA dataset (see results), we then inferred multilocus coalescent-based Extended Bayesian Skyline Plots (EBSP; Heled and Drummond 2008) implemented in BEAST v2.7.6 (Bouckaert et al. 2019) to estimate changes in effective population sizes over time. To infer the demographic history using the nuDNA dataset, we first generated a folded SFS file for the SNP dataset with easySFS (Gutenkunst et al. 2009). We then used Stairway Plot v. 2.18 (Liu and Fu 2020) to estimate historical effective population size changes of *Tropidurus xanthochilus*, *T. tarara*, *T. spinulosus*, and *T. guarani*. At this point, it is important to note that although EBSP, Stairway Plot, and Tajima’s D were applied to monophyletic clusters below the species level, these methods are designed for unstructured samples drawn from local populations. Consequently, the presence of intraspecific population structure violates their underlying assumptions. In coalescent-based approaches, for instance, such structure often results in inflated estimates of historical effective population size (e.g., Heller et al. 2013). While we proceeded with these analyses, we acknowledge that their results may be biased and interpret them with extreme caution. Please refer to the Supplementary Material, section “Details on demographic inference methodology and results”, for a detailed explanation of these tests.

Table 1 provides a summary of all analyses conducted in this study, along with their respective objectives and main results.

**Table 1.**
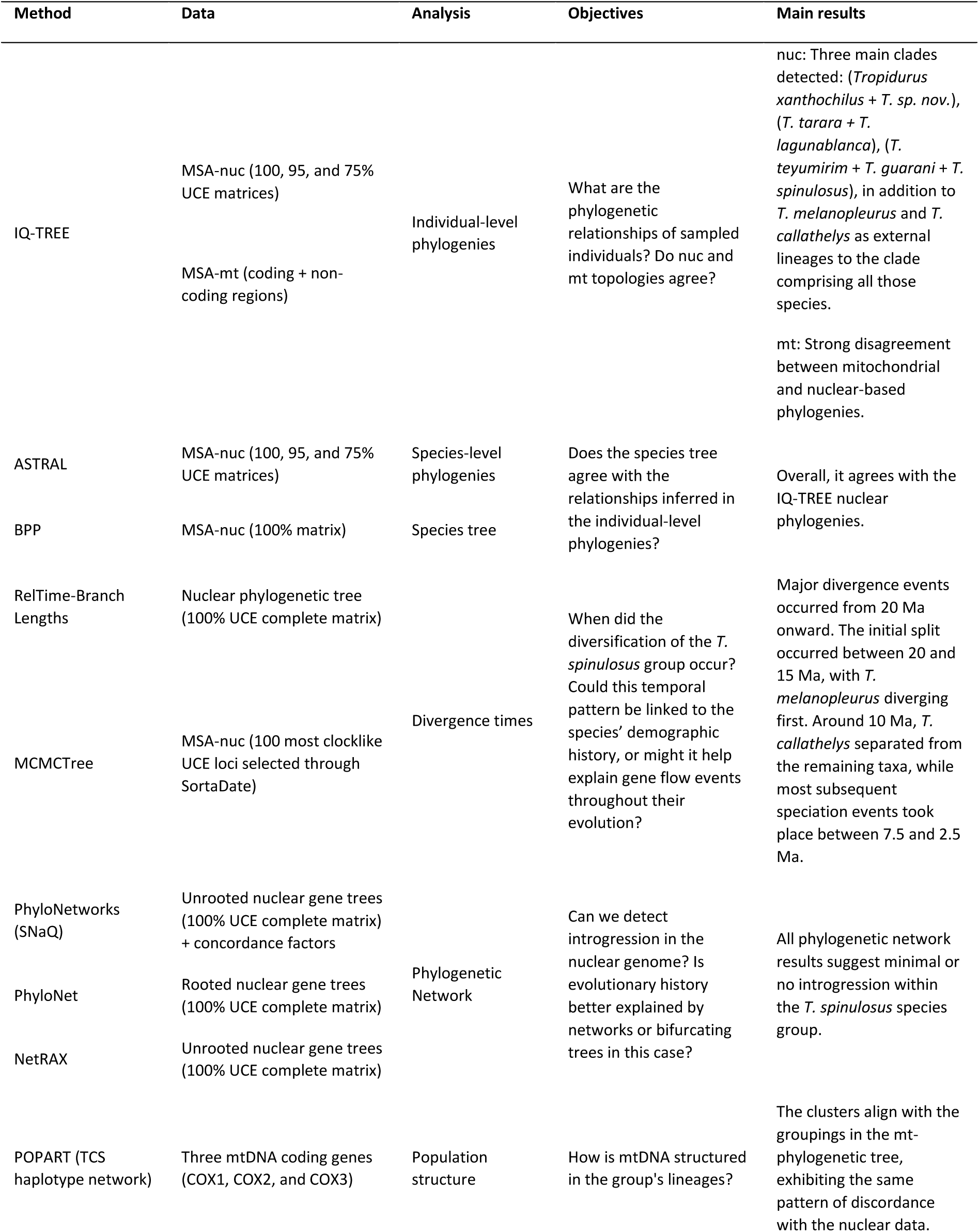
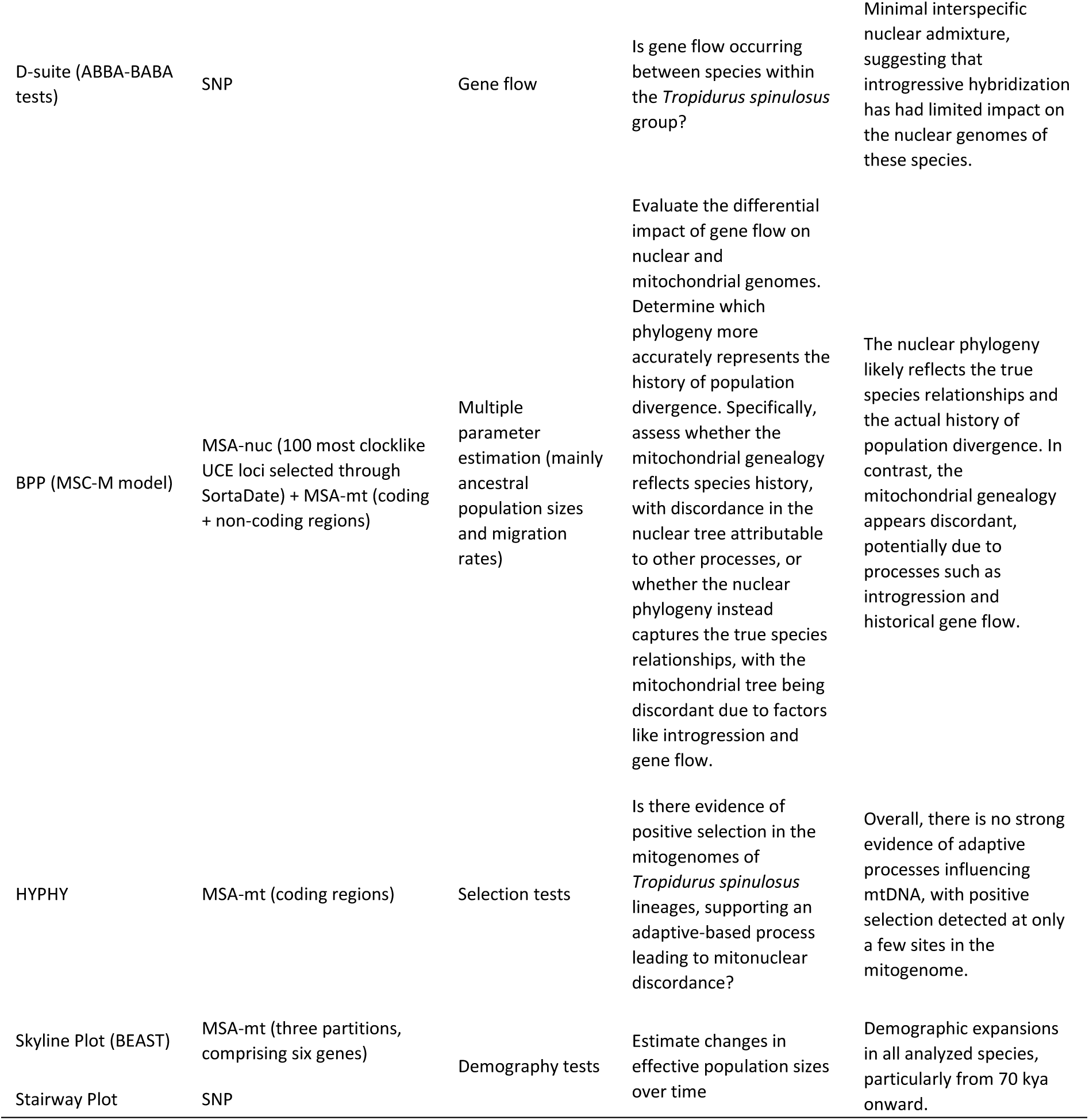
Overview of the analytical workflow used in this study, detailing the rationale behind each step. SNP = single nucleotide polymorphism; MSA = multiple sequence alignment; nuc = nuclear; mt = mitochondrial.

| Method | Data | Analysis | Objectives | Main results |
| --- | --- | --- | --- | --- |
| IQ-TREE | MSA-nuc (100, 95, and 75% UCE matrices) | Individual-level phylogenies | What are the phylogenetic relationships of sampled individuals? Do nuc and mt topologies agree? | nuc: Three main clades detected: (*Tropidurus xanthochilus* + *T. sp. nov.*), (*T. tarara* + *T. lagunablanca*), (*T. teyumirim* + *T. guarani* + *T. spinulosus*), in addition to *T. melanopleurus* and *T. callathelys* as external lineages to the clade comprising all those species. |
|  | MSA-mt (coding + non-coding regions) |  |  | mt: Strong disagreement between mitochondrial and nuclear-based phylogenies. |
| ASTRAL | MSA-nuc (100, 95, and 75% UCE matrices) | Species-level phylogenies | Does the species tree agree with the relationships inferred in the individual-level phylogenies? | Overall, it agrees with the IQ-TREE nuclear phylogenies. |
| BPP | MSA-nuc (100% matrix) | Species tree |  |  |
| RelTime-Branch Lengths | Nuclear phylogenetic tree (100% UCE complete matrix) | Divergence times | When did the diversification of the *T. spinulosus* group occur? Could this temporal pattern be linked to the species' demographic history, or might it help explain gene flow events throughout their evolution? | Major divergence events occurred from 20 Ma onward. The initial split occurred between 20 and 15 Ma, with *T. melanopleurus* diverging first. Around 10 Ma, *T. callathelys* separated from the remaining taxa, while most subsequent speciation events took place between 7.5 and 2.5 Ma. |
| MCMCTree | MSA-nuc (100 most clocklike UCE loci selected through SortaDate) |  |  |  |
| PhyloNetworks (SNaQ) | Unrooted nuclear gene trees (100% UCE complete matrix) + concordance factors | Phylogenetic Network | Can we detect introgression in the nuclear genome? Is evolutionary history better explained by networks or bifurcating trees in this case? | All phylogenetic network results suggest minimal or no introgression within the *T. spinulosus* species group. |
| PhyloNet | Rooted nuclear gene trees (100% UCE complete matrix) |  |  |  |
| NetRAX | Unrooted nuclear gene trees (100% UCE complete matrix) |  |  |  |
| POPART (TCS haplotype network) | Three mtDNA coding genes (COX1, COX2, and COX3) | Population structure | How is mtDNA structured in the group's lineages? | The clusters align with the groupings in the mt-phylogenetic tree, exhibiting the same pattern of discordance with the nuclear data. |
| D-suite (ABBA-BABA tests) | SNP | Gene flow | Is gene flow occurring between species within the *Tropidurus spinulosus* group? | Minimal interspecific nuclear admixture, suggesting that introgressive hybridization has had limited impact on the nuclear genomes of these species. |
| BPP (MSC-M model) | MSA-nuc (100 most clocklike UCE loci selected through SortaDate) + MSA-mt (coding + non-coding regions) | Multiple parameter estimation (mainly ancestral population sizes and migration rates) | Evaluate the differential impact of gene flow on nuclear and mitochondrial genomes. Determine which phylogeny more accurately represents the history of population divergence. Specifically, assess whether the mitochondrial genealogy reflects species history, with discordance in the nuclear tree attributable to other processes, or whether the nuclear phylogeny instead captures the true species relationships, with the mitochondrial tree being discordant due to factors like introgression and gene flow. | The nuclear phylogeny likely reflects the true species relationships and the actual history of population divergence. In contrast, the mitochondrial genealogy appears discordant, potentially due to processes such as introgression and historical gene flow. |
| HYPHY | MSA-mt (coding regions) | Selection tests | Is there evidence of positive selection in the mitogenomes of *Tropidurus spinulosus* lineages, supporting an adaptive-based process leading to mitonuclear discordance? | Overall, there is no strong evidence of adaptive processes influencing mtDNA, with positive selection detected at only a few sites in the mitogenome. |
| Skyline Plot (BEAST) | MSA-mt (three partitions, comprising six genes) | Demography tests | Estimate changes in effective population sizes over time | Demographic expansions in all analyzed species, particularly from 70 kya onward. |
| Stairway Plot | SNP |  |  |  |

## Results

### UCE and mtDNA phylogenomic datasets

We obtained a final dataset including 820 UCE loci for 43 individuals of nine lineages of the *Tropidurus spinulosus* species group. Our complete matrix comprised alignments with 725,505 bp after internal-trimming. Matrices with 75% and 95% of completeness generated alignments with 2,269 and 1,613 loci, 1.98M and 1.40M bp, respectively. Nuclear phylogenetic analyses using both the partitioned approach in IQ-TREE, based on complete and incomplete matrices (Figures S1-S3), and the BPP species tree approach (Figure 2), indicated that our samples can be grouped into three main clades: (*Tropidurus* sp. nov. *+ T. xanthochilus), (T. lagunablanca + T. tarara), (T. guarani + T. spinulosus + T. teyumirim)*, in addition to *T. callathelys and T. melanopleurus* as external lineages to the clade comprising the remaining species. The trees inferred using ASTRAL, which account for ILS, produced similar patterns (Figures S4-S6). Compared to the UCE dataset, the mtDNA dataset (alignment with 40 samples, 15 genes, and 13,578 bp) produced markedly different results regarding topology, indicating a clear case of mitonuclear discordance (Figure 2; Figures S7-S8). Notably different from the nuclear trees, our mitochondrial genealogy shows that while most *T. xanthochilus* samples were grouped as sister to the clade containing *T. spinulosus* and *T. guarani*, two *T. xanthochilus* samples, the *T. sp. nov.* and the *T. teyumirim* samples, were all recovered within the *T. tarara* + *T. lagunablanca* clade. Furthermore, despite *T. xanthochilus* and *T. sp. nov.* forming a clade based on nuclear data, they do not appear closely related in the mitochondrial dataset. Similarly, most groups observed in these two phylogenetic trees are reflected in the haplotype network inferred for the *T. spinulosus* group, showing the same pattern of discordance compared to the nuclear data (Figure S9).

**Figure 2.**
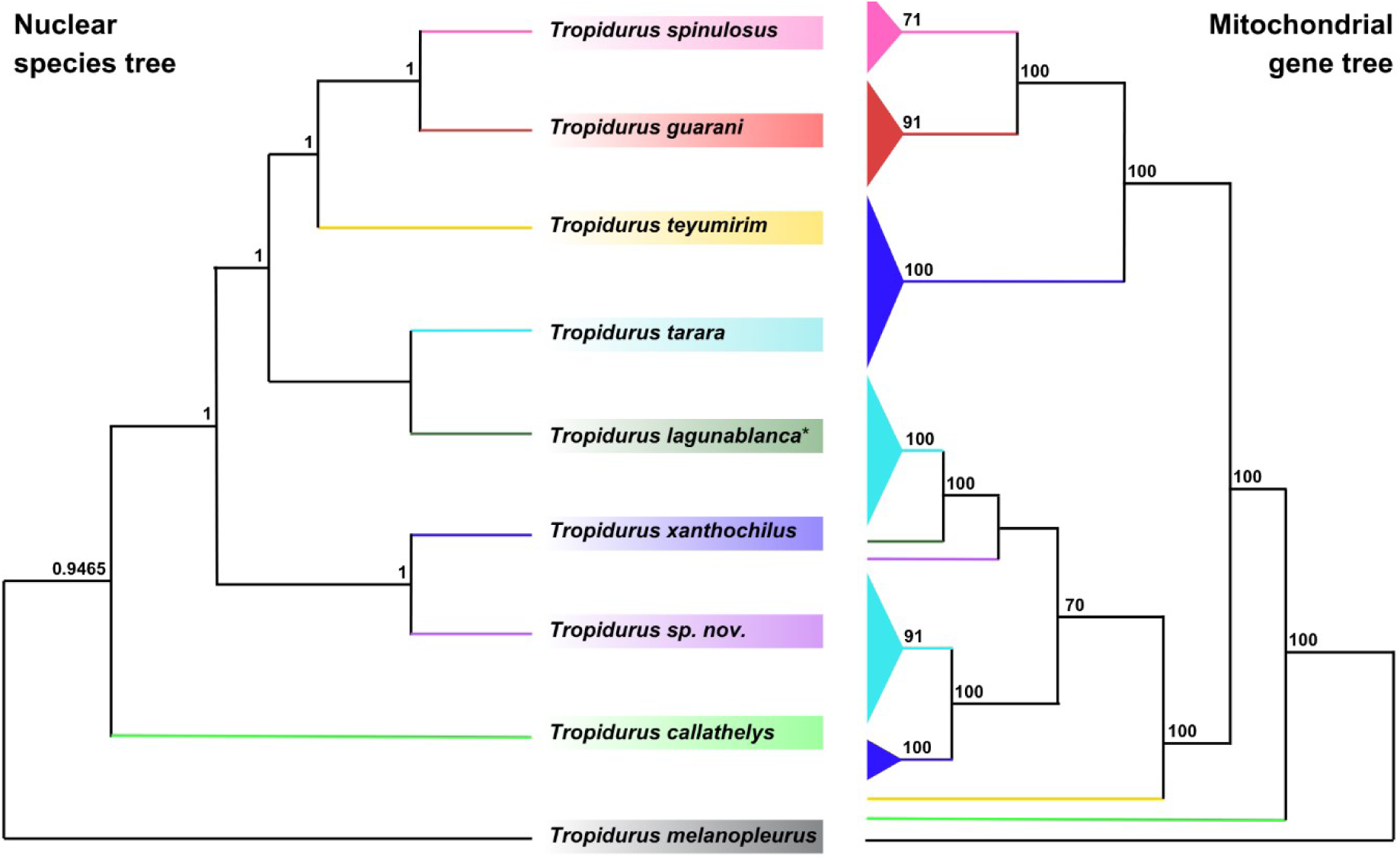
Comparison between the nuclear species tree and the mitochondrial gene tree for the group *Tropidurus spinulosus*. The left panel shows the species tree inferred from multilocus nuclear data using BPP, with branch support values indicated next to nodes (posterior probabilities). The right panel displays the mitochondrial gene tree inferred through IQ-TREE, with branch support shown as bootstrap values (in this case, we only show values above 70). Colored clades correspond to the same taxa in both trees. Collapsed branches in the mitochondrial tree represent intraspecific variation (for the fully detailed tree, see the Supplementary Material). *All nuclear gene trees recovered *T. lagunablanca* nested within *T. tarara* samples, indicating that these lineages likely represent a single evolutionary unit. Accordingly, the *T. lagunablanca* sample was assigned to *T. tarara* in the BPP analysis, so that they represent a single branch in the estimated species tree. However, for a clearer comparison with the mitochondrial genealogy, here we present the two lineages as separate branches within the same clade.

Divergence dating with nuclear data indicates that the initial split within the *Tropidurus spinulosus* group occurred between 20 and 15 million years ago (Ma), separating *T. melanopleurus* from the other lineages. The split involving *T. callathely*s occurred approximately 10 Ma ago, and most subsequent divergence events among the remaining species of the group took place between 7.5 and 2.5 Ma (Figure 3; Figure S10). In most cases, the divergence times estimated using RelTime fall within the same 95% highest posterior density (HPD) intervals provided by MCMCTree, but some differences can be noticed (Figures S10-11).

**Figure 3.**
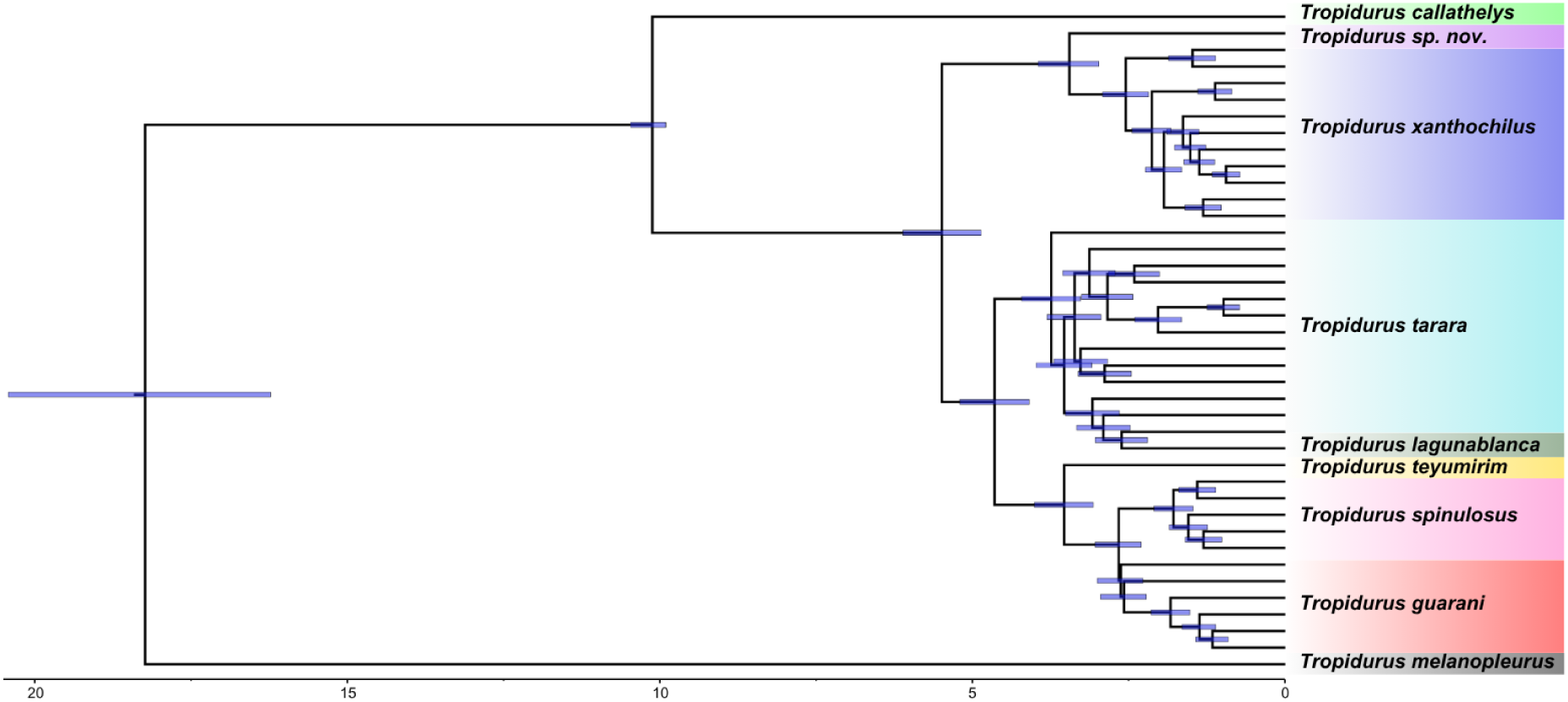
Divergence times (Ma) estimated for the *Tropidurus spinulosus* species group using the UCE dataset. The 95% HPD interval of the posterior estimates (blue shaded bars), as estimated through the MCMCTree program, is shown above each node of the tree. The scale bar is in Ma. Specimen names are collapsed (for the fully detailed tree, see the Supplementary Material).

### Phylogenetic networks and gene flow

Analyses of the UCE dataset using PhyloNetworks revealed that models incorporating up to three reticulation events provided a better fit to our data compared to strictly bifurcating trees, where gene tree discordance is attributed solely to ILS. However, these events are mostly intraspecific and do not involve the species with discordant relationships; thus, Phylonetworks does not indicate that reticulation is the primary source of mitonuclear discordance (Figures S12-S15; Table S1). Analyses using the other phylogenetic network approaches corroborated the pattern of minimal reticulations between species observed in our UCE data (Figures S16-S18), and ABBA-BABA tests for detecting gene flow also do not indicate significant nuclear admixture within the *Tropidurus spinulosus* species group (Table S2). Please refer to the Supplementary Material, section “*Details on Phylogenetic Networks methodology and results*,” for a detailed explanation of all reticulation tests and respective results.

Analyses in BPP under the MSC-M model with nuclear data indicate minimal or no gene flow among extant species, but inclusion of ancestral populations yields higher migration rate estimates (Figure 4; Tables S3-S4), particularly those associated with the observed phylogenetic discordance. This pattern is robust to the choice of different guide trees: whether constrained by the mitochondrial topology or by the partitioned nuclear topology, although the former produces relatively higher migration rate values. Notably, both guide tree inferences implicate *Tropidurus xanthochilus*, or its ancestors, in these ancient gene flow events, suggesting that historical mitochondrial capture involving this lineage may underlie the main observed discordances between nuclear and mitochondrial phylogenies. Also, when combining the nuclear dataset alongside the complete mitochondrial genome, higher rate estimates were generally produced than those obtained using the nuclear dataset only. In this case, migration rates using the combined dataset (with mitochondrial and nuclear partitions) were higher than the estimates using solely the nuclear data (Table S4). In summary, all migration scenarios evaluated, including information derived from the mitochondrial data (whether based on topology or mitogenome alignments), consistently resulted in higher inferred migration rates compared to nuclear-only analyses, demonstrating that the inclusion of mtDNA in the analysis strengthens the detection of historical gene flow in this system.

**Figure 4.**
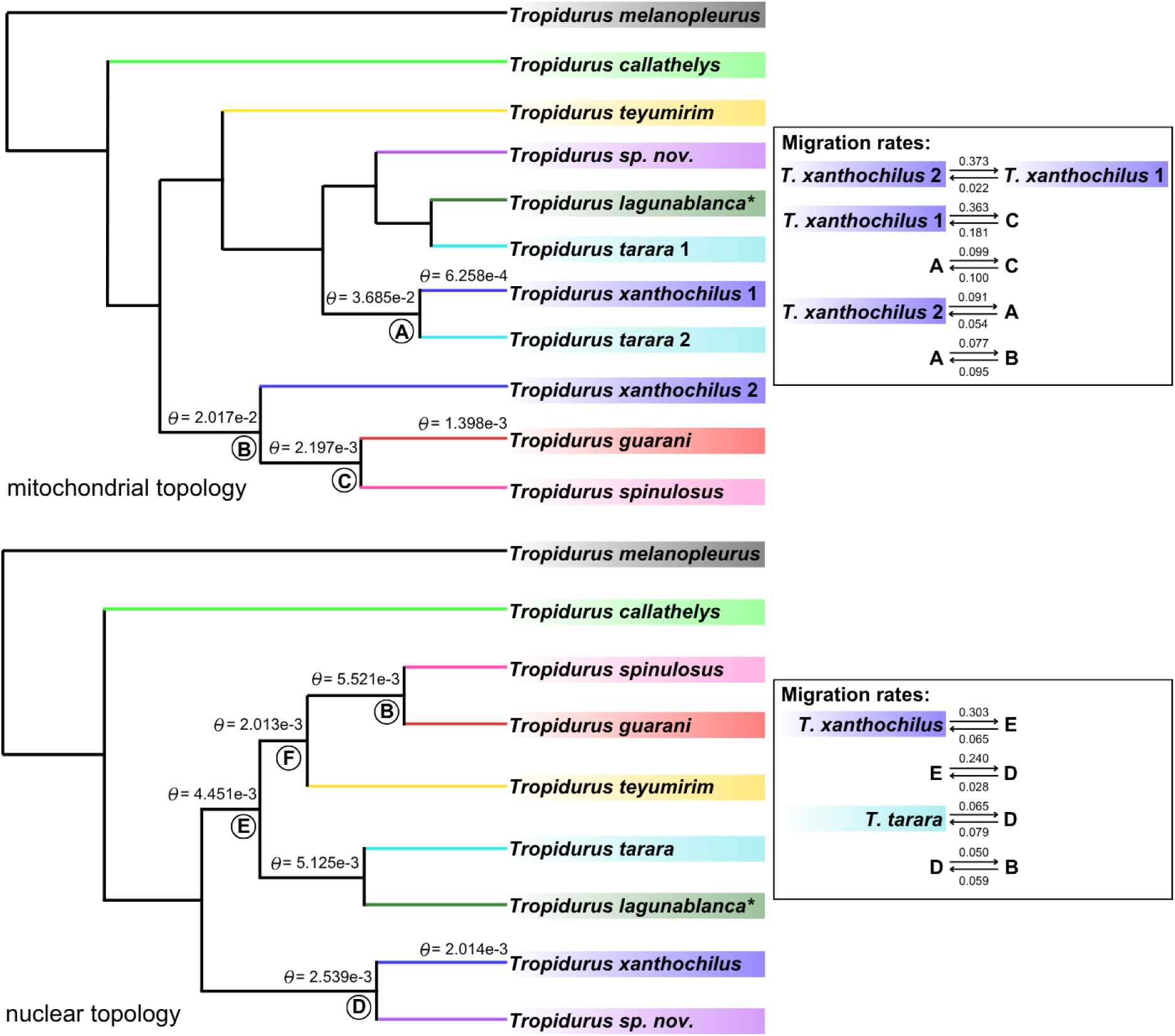
Migration (MSC-M) model for *Tropidurus spinulosus* species, displaying only the scenarios with the highest estimated migration rates. For a complete list of all tested migration scenarios and their corresponding rate values, please refer to the Supplementary Material. Population size (theta, θ) values for each species or ancestral node involved in gene flow events are shown above the branches, with the remaining theta values also provided in the Supplementary Material. The upper panel presents the analysis using the mitochondrial topology as the guide tree; the lower panel shows the same analysis based on the nuclear topology, in this case using both nuclear and mitochondrial data as input. In both trees, the circled letters below the nodes denote the ancestral lineages included in the migration tests.

Estimates of population size (θ) for both current species and ancestral populations further support the previously described scenario (Figure 4; Tables S5-S6), for instance, indicating a directional gene flow event in which the larger ancestral population of the *T. guarani* + *T. spinulosus* clade might have captured *T. xanthochilus* mitochondrial genomes.

### Selection

Regarding the mtDNA dataset, our analyses indicate minimal evidence of positive selection acting on the mitogenomes of *Tropidurus spinulosus* lineages. Codon-based tests detected limited signatures of episodic positive or diversifying selection: MEME identified nine sites (two in ATP6, one in ATP8, one in COX1, two in COX2, and one in COX3) with p-values ≤ 0.05, while FUBAR detected two sites (one in COX1 and one in COX2) with posterior probabilities ≥ 0.95. In contrast, both BUSTED and aBSREL analyses did not reveal significant episodic selection along any branches in the mitogenomic phylogeny, even after formal testing of 36 branches with correction for multiple comparisons. Furthermore, we did not observe any species with extreme nucleotide diversity (π) values, which could indicate selection on mitochondrial lineages; instead, the diversity values were nearly identical across all populations (Table S7). Notably, the significantly negative Tajima’s D (p ≤ 0.05, Table S8) indicates an excess of rare variants, a pattern that can arise from both demographic expansion and selection. Subsequent analyses support a scenario of recent expansion for the main populations, suggesting that this signal is evidence of historical demographic changes rather than selection acting on the mitogenomes.

### Demographic history

As previously demonstrated, BPP estimates of population size reveal differences among ancestral populations likely involved in the ancient introgression events underlying the observed discordance; these differences also support the inferred direction of gene flow (please refer to the *Migration tests* in the *BPP* section of the Supplementary Material as well). Furthermore, EBSP results indicated a marked increase in mitochondrial effective population size across all populations, particularly within the last 100,000 years (Figure S19). In the nuclear dataset, Stairway Plot analyses showed a consistent pattern of population expansion beginning around 60,000 years ago, followed by relatively stable sizes thereafter (Figure S20).

## Discussion

Our genomic analysis of the *Tropidurus spinulosus* species group reveals a complex evolutionary history characterized by: (a) significant discordance between nuclear and mitochondrial phylogenetic groupings, with very low levels of nuclear introgression; (b) at least two major mitochondrial capture events, specifically involving ancient *T. xanthochilus* populations and non-sister groups such as the *T. spinulosus* + *T. guarani* clade and *T. tarara*; (c) no signs of strong positive selection acting on mtDNA; and (d) demographic evidence of recent population expansions in both nuclear and mitochondrial genomes across various species in the group, along with differences in the sizes of ancestral populations that may have reinforced directionality of introgression events. Given these findings, the mitonuclear discordance observed in this study likely originated from admixture among ancient populations of the *T. spinulosus* species group that experienced long-term isolation, followed by secondary contact.

### Methodological challenges in investigating introgression as a cause of mitonuclear discordance

Some caution is warranted when interpreting our results, especially given the nature of our dataset. UCEs, by definition, represent highly conserved genomic regions shared across evolutionarily distant taxa (Faircloth et al. 2012). While this might suggest that such data are primarily suited for addressing ’deep’ phylogenetic questions, multiple studies have demonstrated their utility at the population and individual levels (Smith et al. 2014; Raposo do Amaral et al. 2018). Additionally, the introgression and gene flow tests we used can capture events on different time scales. For instance, the ABBA-BABA test assumes that multiple substitutions at a given site are rare, as an excess of substitutions can distort patterns of site discordance. This assumption tends to break down in deeply divergent taxa (Hibbins and Hahn 2022), making the ABBA-BABA test more suitable for detecting recent introgression events. We detected only one significant case of interspecific gene flow (both in the case of the PhyloNetworks analysis and the ABBA-BABA tests; however, it warrants caution (as discussed below). Conversely, phylogenetic networks, which analyze discordance between gene trees and other concordant factors, are less impacted by multiple substitutions at individual sites, making them better suited for detecting older introgression events (Hibbins and Hahn 2022). Therefore, as we used different methodologies capable of detecting signals across distinct time scales, our results cannot be considered dependent on the nature of the used markers, even considering that the accuracy of phylogenetic network methods may also depend on the number of gene trees used. In this regard, previous studies found that a similar number of gene trees compared to our dataset is sufficient to reliably recover an accurate network topology, even though the exact direction of reticulation events may remain uncertain depending on the evolutionary scenario (Solís-Lemus and Ané 2016). Moreover, the consistency of recovered topologies across multiple analyses, with varying levels of data completeness, suggests that the majority of our UCE gene trees are resolved and phylogenetically informative. Therefore, although other genomic markers may offer greater sensitivity for detecting introgression, the consistency of our results across multiple analytical frameworks—each suited to detecting gene flow over different temporal scales—supports the conclusion of minimal or no nuclear introgression within the *Tropidurus spinulosus* group. In any case, future comparative studies are warranted to evaluate how the choice of genomic markers may influence the detection of complex evolutionary processes such as introgression.

### Decoupled histories: mitonuclear discordance with minimal nuclear introgression

Reports of mitonuclear discordance are not unexpected, given that mitochondrial and nuclear genomes evolve semi-independently. In this context, while previous studies have documented nuclear introgression associated with mitonuclear discordance (e.g., Zieliński et al. 2013; Myers et al. 2022), our findings point to a different scenario. We found little evidence of substantial nuclear genome introgression, and even the two specific results that could indicate historical hybridization within the group—a significant nuclear reticulation event involving *Tropidurus callathelys* (Figure S14) and a significant D-statistic involving *T. teyumirim* (Table S2)—should be interpreted with caution. First, there is no evidence suggesting contemporary hybridization within the *T. spinulosus* group. The known biology and distribution of these specific lineages do not suggest the occurrence of hybridization events between them (Carvalho et al. 2013; Carvalho 2013; 2016), although this may reflect a distinction between contemporary gene flow and the ability of our methods to detect introgression over evolutionary timescales. Second, the extreme sampling disparity (n=1 for both *T. teyumirim* and *T. callathelys* versus larger samples for other taxa) could have contributed to false positives in ABBA-BABA tests, which have reduced sensitivity at low admixture levels (Durand et al. 2011), and PhyloNetworks analyses due to stochastic effects.

ILS is another phenomenon that can contribute to mitonuclear discordance, and its signals can easily be mistaken for those caused by introgression (Andersen et al. 2021; DeRaad et al. 2023). In our case, if some samples of a species coalesced deep in the tree, before this lineage diverged from its common ancestor, a significant portion should have coalesced if ILS were the predominant factor driving the discordance with mitochondrial data. However, even our phylogenetic network analyses, which explicitly account for ILS (particularly PhyloNetworks and PhyloNet), did not detect substantial nuclear introgression events. Moreover, the agreement between partitioned (based on IQ-TREE) and ASTRAL analyses of nuclear markers also suggests that ILS has a limited impact on the phylogenetic signal (Figures S1-S6), as ASTRAL is designed to recover the correct topology even under high levels of ILS (Zhang et al. 2018).

Finally, it is essential to acknowledge the lack of methods specifically designed to detect mitochondrial genome introgression with the same level of precision available for nuclear markers. The approach adopted in this study—particularly the BPP-based framework using both mitochondrial and nuclear topologies as guide trees and integrating the mitochondrial partition with multiple nuclear loci (Figure 4)—was designed to illustrate how gene flow may differ between nuclear and mitochondrial genomes due to their distinct inheritance patterns and evolutionary dynamics. Furthermore, we sought to highlight the value of a genome-wide analytical perspective for elucidating the role of introgressive hybridization in animal speciation—an approach still underrepresented in many biological systems (Good et al. 2015). Notably, recent studies have shown that, even in contact zones with frequent hybridization, genetic exchange can remain confined to very narrow geographic ranges, indicating that mitochondrial transfer does not necessarily lead to widespread introgression (Zozaya et al. 2024). This contrast illustrates how the relative permeability of nuclear and mitochondrial genomes to gene flow can vary markedly among squamate clades, underscoring the utility of our BPP-based framework for revealing a distinct pattern in *Tropidurus*.

Specifically, we discovered that while our nuclear data largely dismisses the occurrence of prominent introgression within our species group, the mitochondrial genome presents a contrasting picture. Migration rate estimates increase when mitochondrial DNA is included— either as an additional partition or by using the mitochondrial topology as the guide tree (Tables S3-S4). Specifically, while BPP analyses of 100 UCE loci produced minimal migration rate estimates (under the nuclear-guided topology), incorporating the mitogenome into the dataset increased the estimated values despite the much larger size of the nuclear dataset. Notably, higher migration rate estimates appeared only in scenarios involving ancestral populations (regardless of whether the nuclear or mitochondrial topology was used as the guide tree), producing more realistic values (Figure 4). These findings not only underscore the analytical value of including mitochondrial data in gene flow estimates but also support the conclusion that the nuclear phylogenies more accurately reflect species history in the *T. spinulosus* system.

### Proximate explanations of mitonuclear discordance in the Tropidurus spinulosus species group

Evidence supporting a demography-related hypothesis regarding introgression typically falls into three main categories (e.g., Hinojosa et al. 2019; Palacios et al. 2023; Shen et al. 2025): (i) prolonged periods of isolation; (ii) population growth leading to secondary contact; and (iii) a biogeographic signal related to the discordance. Below, we discuss how our findings align with this framework, as the primary cause of the mitonuclear discordance reported in this study is mitochondrial capture associated with the demography of the lineages within the *Tropidurus spinulosus* group. Specifically, our results indicate ancient gene flow events occurring in populations that were likely isolated for extended periods before undergoing secondary contact, although we detected limited evidence of nuclear introgression. Additionally, both nuclear and mitochondrial datasets reveal widespread historical fluctuations in population size (Figures S19-S20; Tables S5-S6). Similar cases have been documented in lizards (e.g., Myers et al. 2022), where geographic isolation, followed by range and demographic expansions, led to secondary contact and gene flow between previously diverged groups.

First, our divergence time estimates support a gradual and prolonged separation among species in the *Tropidurus spinulosus* group, with divergence events between species occurring from 20 Ma onward. Specifically, the initial split occurred between 20 and 15 Ma, with *T. melanopleurus* diverging first. Around 10 Ma, *T. callathelys* separated from the remaining taxa, while most subsequent speciation events took place between 8 and 2.5 Ma (Figure 3). Related to that, it is essential to note that the evolutionary history of South America’s dry diagonal has been shaped by multiple Neogene (23 Ma onward) geological and climatic processes, with individual taxa responding uniquely to local conditions (Guillory et al. 2024). Deep divergences among Squamates, including those within the *T. spinulosus* group, likely stem from Miocene events such as the uplift of the Central Brazilian Plateau and marine incursions in the southern dry diagonal (Guillory et al., 2024; Bezerra et al., 2025).

If such geological events led to prolonged geographic isolation of lineages, their gradual cessation would have increased opportunities for secondary contact and potential introgression, particularly when coupled with demographic and range expansions. In such a scenario, periods of secondary contact may have led to the formation of hybrid zones with varying levels of interbreeding, followed by contrasting patterns of gene flow between the nuclear and mitochondrial genomes. During periods of geographic and/or demographic expansion, nuclear genomes—subject to recombination and meiotic segregation—undergo widespread admixture and tend to homogenize over time, diluting distinct traces of past isolation and secondary contact. In contrast, divergent mitochondrial haplotypes, which are non-recombining and inherited solely from the females, may persist due to genetic drift, particularly if introgression occurred early in the population expansion or when the population sizes were relatively low.

The directionality of gene flow events is another factor that could have played a crucial role in facilitating neutral mitochondrial capture. Because genetic drift is more pronounced in small populations, neutral mutations in mitogenomes are more likely to become fixed (Moore 1995; Després, 2019). Consequently, unidirectional introgression from a larger, resident population into a smaller, invading one would be more probable during the early stages of contact. If the ancestral populations of any species in the *Tropidurus spinulosus* group were more range-restricted, such conditions would further facilitate the fixation of captured mitochondrial variants through drift. The repeated instances of discordance involving *T. xanthochilus*, due to ancient introgression, may suggest that this lineage had a smaller population size compared to others. Our ancestral population size estimates are consistent with this scenario. For instance, in the model where migration rates are higher (in this case, using the mitochondrial topology as the guide tree; Tables S3-S4), Θ estimates of *T. xanthochilus* are considerably lower than those of the ancestor involved in the gene flow event—in this case M (*T. xanthochilus* ^͢^ ancestor (*T. guarani*, *T. spinulosus*)) = 0.3629, with Θ values of 0.0006258 and 0.02017, respectively (Table S5). A similar pattern was observed when the nuclear topology was used as the guide tree, with the ancestors of the *T. tarara* + *T. spinulosus* + *T. guarani* clade exhibiting higher Θ values than those of the ancestral *T. xanthochilus* populations (Table S6). Under this scenario, it would be plausible to suggest that the mitogenomes of some *T. xanthochilus* individuals could have been incorporated by the larger ancestral populations of the *T. guarani* + *T. spinulosus* clade, and also by ancestral populations of *T. tarara*.

It is also noteworthy to emphasize that two of our demographic analyses (EBSP and Stairway Plot) may capture only a narrower temporal window than the full span of historical gene-flow events. Furthermore, both approaches assume panmictic, unstructured populations (an assumption likely violated by our sampling of divergent lineages), which can bias demographic reconstructions (Heller et al. 2013). Even so, it is reasonable to assume that relatively recent demographic shifts during the Pleistocene (2.5 Ma–12 ka) may have contributed to the observed mitonuclear discordance, either by facilitating the fixation of mitochondrial variants or by reducing detectable signatures of nuclear introgression. Our results suggest that population expansions might have begun at least 100 ka ago in the mitochondrial dataset (Figure S19) and around 60 ka in the nuclear dataset (Figure S20), with relative stability thereafter. Similar late-Pleistocene demographic fluctuations shaping phylogeographic patterns have been documented in other squamate lineages (Guillory et al. 2024). Finally, the temporal mismatch between mitochondrial and nuclear estimates likely reflects the distinct mutation rates applied to each dataset; as with many non-model organisms, specific substitution rate estimates for *Tropidurus* species are unavailable, adding uncertainty to absolute time estimates. For these reasons, we interpret our demographic inferences with caution, recognizing that both population structure and rate uncertainty may influence the magnitude and timing of inferred events.

In addition, the two particular instances of mtDNA capture involving *Tropidurus xanthochilus* are also associated with biogeographic signals: one possibly occurring to the west of *T. xanthochilus*’ distribution, involving ancestral populations of the *T. spinulosus + T. guarani* clade, and another to the south, involving *T. tarara*. *Tropidurus spinulosus* is found west of the Paraguay River, from north-central Argentina and northwestern Paraguay to southeastern and central Bolivia, covering both the Chaco and dry forest zones (Frost et al. 1998; Carvalho 2013). Although distributional data remain limited, the known ranges of *T. spinulosus* and *T. xanthochilus* are allopatric. The nearest known populations of *T. spinulosus* are located at least 350 km south of the type locality of *T. xanthochilus,* near the ‘*Serranía de Huanchaca*’ (Harvey and Gutberlet, 1998). Even so, Harvey and Gutberlet (1998) proposed that *T. xanthochilus* and *T. spinulosus* might be parapatric in the area where the forests of the Tarvo and Paraguá Rivers merge with the semideciduous *‘Chiquitania’* dry forest. On the other hand, *T. tarara* is known from the northern portion of Eastern Paraguay, east of the Paraguay River, to the Brazilian state of Mato Grosso do Sul (near the southern limit of the distribution of *T. xanthochilus*).

Future studies (whether empirical or based on modelling scenarios) will be crucial to assess the potential overlap (past or present) in the distributions of *Tropidurus xanthochilus* with *T. spinulosus* + *T. guarani* clade, and *T. tarara* as well. Such overlap would provide additional biogeographical evidence for mitochondrial capture among these lineages. Environmental changes can increase opportunities for hybridization and introgression through neutral demographic processes (Excoffier et al. 2009; Phuong et al. 2016), with mitonuclear discordance of this nature often linked to climatic instability and its effects on demographic dynamics (Phuong et al. 2016). Phylogenetic niche modeling, which combines past and present climate data with occurrence records and time-calibrated phylogenies to reconstruct historical geographic ranges (e.g., Guillory and Brown, 2021), may be beneficial for testing these hypotheses in the future.

Finally, our findings suggest that mitochondrial capture in the *Tropidurus spinulosus* group likely occurred during a period when the current lineages were not yet fully differentiated and still overlapped in both space and time. This interpretation is supported by our migration rate results, which show that the highest estimated values were associated with scenarios involving ancestral populations (Figure 4). Notably, this aligns with the “gray zone of speciation” concept—a transitional phase where populations shift from widespread admixture to complete genetic isolation (Roux et al. 2016; Burbrink et al. 2021), facilitated by pre- and/or post-zygotic barriers (Stankowski and Ravinet, 2021). The complexity of our evolutionary scenario also suggests that a bibliographic synthesis indicating the major evolutionary drivers and consequences of introgression and genomic capture across taxonomic groups and biogeographic regions providing research avenues and broad hypothesis-testing frameworks would be of great interest to the evolutionary biology community.

### Mitonuclear discordance beyond neutral processes: a case for natural selection?

Our analyses did not reveal strong evidence of positive selection acting on the mitochondrial genome of the *Tropidurus spinulosus* species group. Site-model tests across the phylogeny identified only a few individual codons under selection, indicating that these methods captured limited signals of episodic or pervasive positive selection acting on the mitogenome. However, these approaches are designed to detect selection at the level of specific codons or phylogenetic branches, rather than to assess the rapid fixation of mitochondrial haplotypes following introgression (see the [HyPhy documentation](https://hyphy.org/methods/general/) for a detailed explanation). In such a scenario, an entire haplotype could be fixed in a new species, a process that might not be directly detectable by the methods adopted here.

Nonetheless, complementary analyses, including nucleotide diversity estimates and Tajima’s D test results, do not provide support for natural selection either. First, comparable π levels across the lineages involved in the discordance (Table S7) are not indicative of selective forces strongly acting on the mitogenomes of *Tropidurus spinulosus* species. Similarly, the negative Tajima’s D values (Table S8), while theoretically compatible with both selection and population expansion, are here interpreted as indicative of historical demographic changes in light of the additional evidence previously discussed. Furthermore, in lizards—where mitochondrial function is less constrained by high metabolic demands compared to endotherms (Chung and Schulte 2020)—the selective pressure for rapid fixation of an introgressed mitogenome is likely lower. Nonetheless, while selective factors are unlikely to be the primary drivers of mitonuclear discordance in *T. spinulosus* lineages, adaptive explanations may still be underrepresented in the literature (e.g., Mao and Rossiter 2020), which may partly reflect the inherent challenges in reliably detecting selection on mtDNA (Bonnet et al. 2017).

### Sex-biased dispersal in Tropiduridae and its potential role in mitonuclear discordance

Finally, future investigations could look further into the role of mitonuclear incompatibilities and mating systems in explaining contrasting signals between nuDNA and mtDNA. Asymmetries between sexes in mating behavior, offspring production, and dispersal are well-known demographic factors contributing to mitonuclear discordance (Dessi et al. 2022). For example, in species where males exhibit greater dispersal, nuDNA is expected to show higher genetic homogeneity across distant populations, as male-mediated gene flow facilitates the spreading and mixing of their genetic material. Conversely, if females are more philopatric (i.e., remain near their birthplace), mtDNA may display greater population structure, preserving historical patterns of isolation. As a result, while nuDNA may indicate a single, cohesive population over a broad geographic range, mtDNA could reveal distinct subpopulations shaped by limited female dispersal. This process can lead to geographically structured mitochondrial lineages that persist despite widespread nuclear gene flow (see reviews in Toews and Brelsford 2012; Bonnet et al. 2017). Currently, empirical data on the dispersal behavior of *Tropidurus* species are lacking. However, among close relatives of Tropiduridae (e.g., Phrynosomatidae), females exhibit strong philopatry and possibly territoriality, whereas males may disperse further from contact zones, facilitating nuclear gene flow but limiting mitochondrial inheritance (Qi et al. 2013). Although limited studies have been conducted on squamate reptiles, some evidence indicates that sex-biased differences in behavior and fitness can influence dispersal patterns. Therefore, we recommend future research to investigate whether sex-related dispersal contributes to the mitonuclear discordance observed in *Tropidurus*.

## Data availability

The data underlying this article, including phylogenetic datasets, corresponding trees, input and output files for all analyses, and any other relevant supplementary files, are available in Zenodo, at https://doi.org/10.5281/zenodo.15376373.

## Author contributions

Matheus Salles: study design, analyses, lead writing. Fabricius Domingos and André Carvalho: study design, field sampling, genome sequencing, writing final draft. Nicolas Martinez, Frederick Bauer, Martha Motte, Viviana Espínola, Miguel T. Rodrigues, Carla Piantoni, Guarino R. Colli, Fernanda P. Werneck: field sampling efforts and/or provision of collection materials and infrastructure. Márcio Pie, André Olivotto, Erik Choueri, Adam Leache: lab work, genome sequencing. All authors read, commented, and approved the final draft of the manuscript.

## Acknowledgments

We thank all the institutions that provided technical, material, and structural support during the sampling phase of this study, namely the Herpetological Collection of the University of Brasília (CHUNB), the Amphibians and Reptiles Collection of the National Institute of Amazonian Research (INPA-H), the Biosciences Institute of the University of São Paulo (IB-USP), the American Museum of Natural History (AMNH), the Museo Nacional de Historia Natural del Paraguay (MNHNP), the Mato Grosso Federal University (UFMT), Museo de Historia Natural Alcide d’Orbigny, and the Universidade Nacional de Salta (UNSa). We extend our gratitude to our esteemed colleagues Eliana Lizarraga, Felipe F. Curcio (UFMT), Karina Atkinson and Jean-Paul Brouard (Fundación Para La Tierra, Paraguay), Johanna López, Nestor Romero (and family) (Estancia Cerrados del Tagatiya), Pastor Enmanuel Pérez-Estigarribia (and family), Ricardo Céspedes, Rodrigo Ayala, and Rolando Rivas for crucial assistance or support during fieldwork. We thank Angela Coda for allowing access to the Reserva Natural Cerrados del Tagatiya and Malvina Duarte for allowing access to the Reserva Natural Laguna Blanca. We also thank Junior Nadaline and Carlos Daniel Rivadeneira for their invaluable assistance with genetic sample processing and DNA sequencing. MTR also thanks all members of his lab for their help in the field. Finally, we extend our gratitude to Prof. Renato Pires Machado (UFPR) for his valuable contributions throughout the corresponding author’s PhD project.

## Benefit-Sharing Statement

A research collaboration was developed with scientists from the countries providing genetic samples; all collaborators are included as co-authors, the research results have been shared with the broader scientific community, and the research addresses a priority concern—the evolutionary history of organisms being studied. More broadly, our group is committed to international scientific partnerships, as well as institutional capacity building.

