## Supplementary for "Ancient introgression explains mitochondrial genome capture and mitonuclear discordance among South American collared *Tropidurus* lizards"

---

---

*Partitioned phylogenetic and divergence time results*.............................................. 2-8
Supplementary Figures S1-S11

*Details on phylogenetic networks methodology and results*.................................. 8-15
Supplementary Figures S12-S18
Supplementary Tables S1-S2

*Migration tests in BPP*........................................................................................ 16-19
Supplementary Tables S3-S6

*Details on demographic inference methodology and results* ................................. 19-22
Supplementary Figures S19-S20

*Remaining supplementary tables*........................................................................ 23-26
Supplementary Tables S7-S10

*Partitioned phylogenetic and divergence time results*

ASTRAL and nuclear gene trees recovered *Tropidurus guarani* as paraphyletic, indicating that this name could either represent a paraphyletic species or a species complex in need of revision (Figures S1-S6). Besides, *T. lagunablanca* appeared nested among samples of *T. tarara*, suggesting that these two lineages likely constitute a single evolutionary entity.

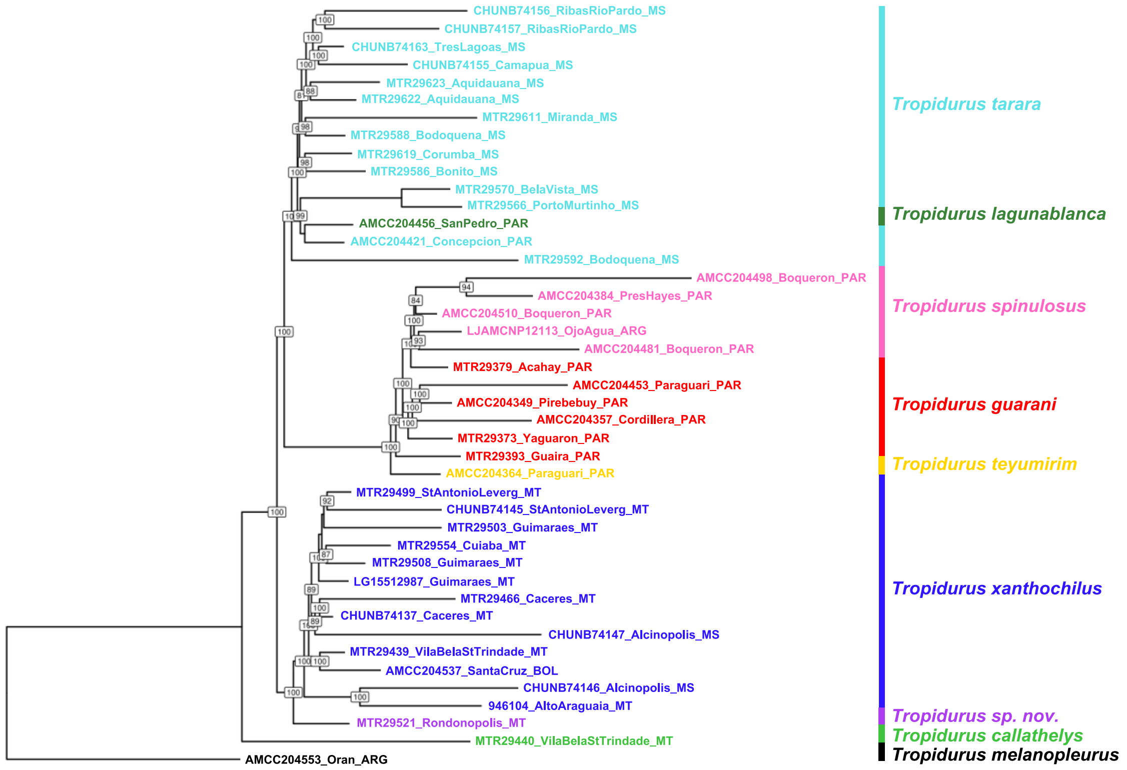


Figure S1. Partitioned phylogenetic analysis of 2269 UCE loci, matrix with 75% of completeness (via IQ-TREE). Numbers inside boxes refer to bootstrap support values (only nodes with support > 80 are shown).

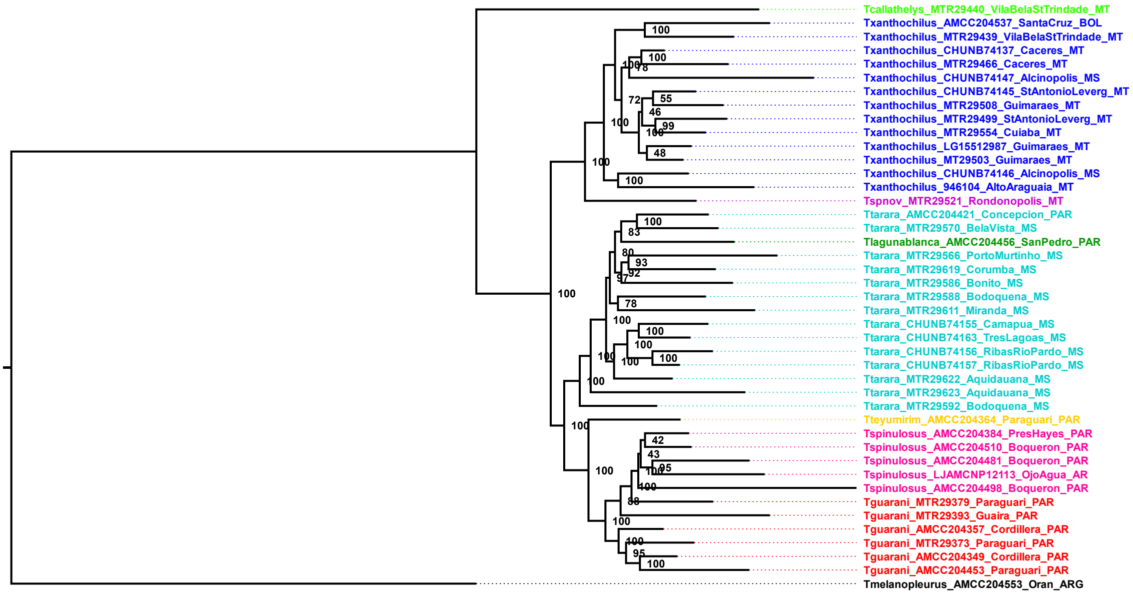


Figure S2. Partitioned phylogenetic analysis of 820 UCE loci (complete matrix), produced via IQ-TREE. Numbers next to nodes refer to bootstrap support values.

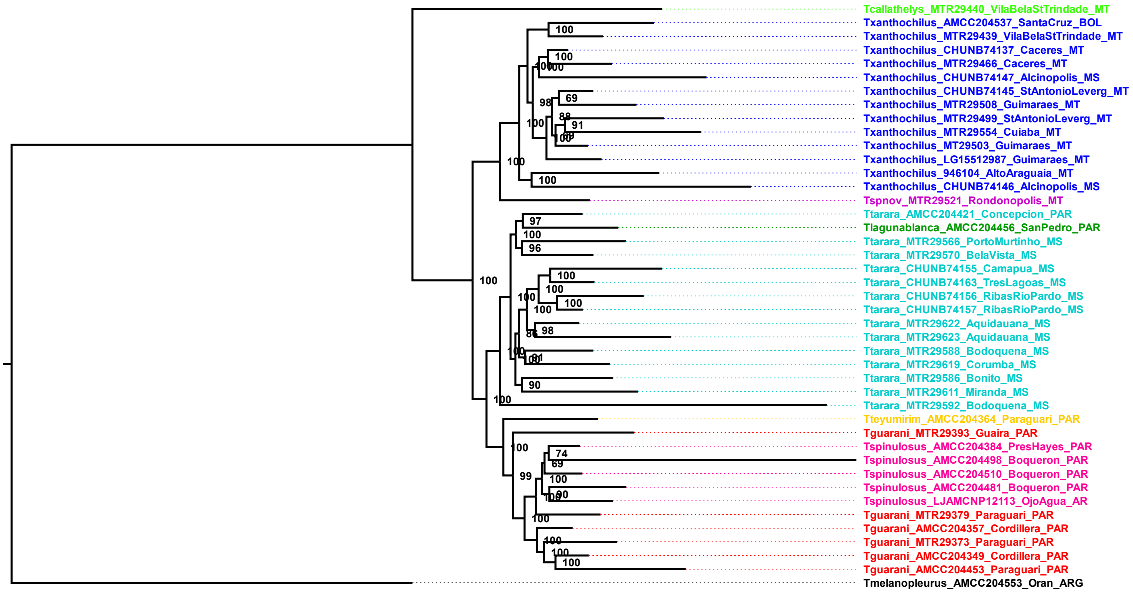


Figure S3. Partitioned phylogenetic analysis of 1613 UCE loci (matrix with 95% of completeness), produced via IQ-TREE. Numbers next to nodes refer to bootstrap support values.

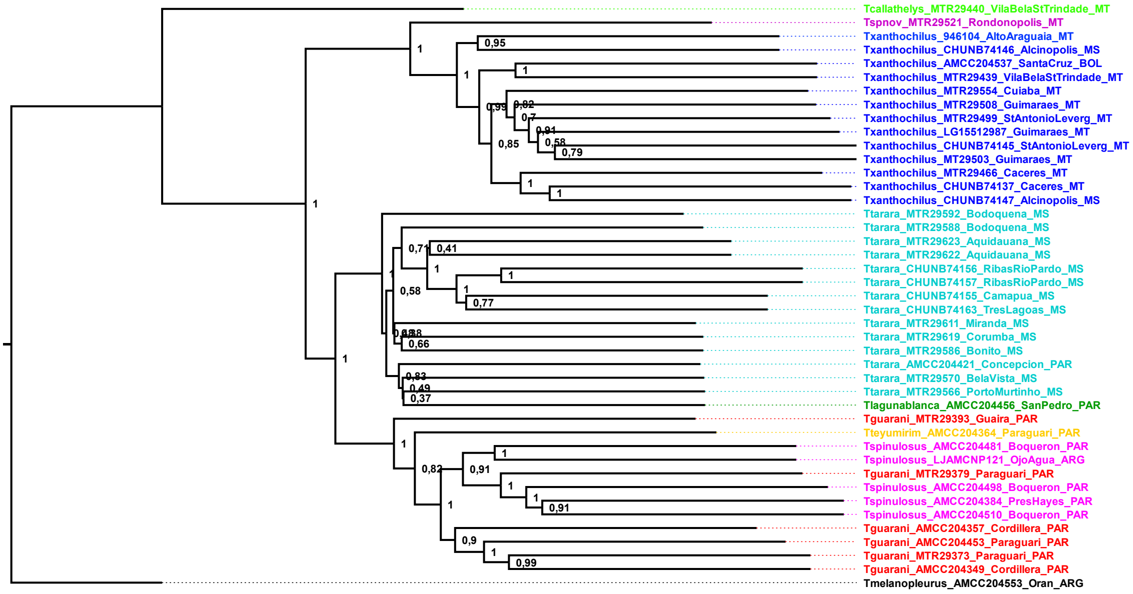


Figure S4. Results produced from the species tree analysis of the 820 UCE loci (complete matrix), produced via ASTRAL. Numbers next to nodes refer to bootstrap support values.

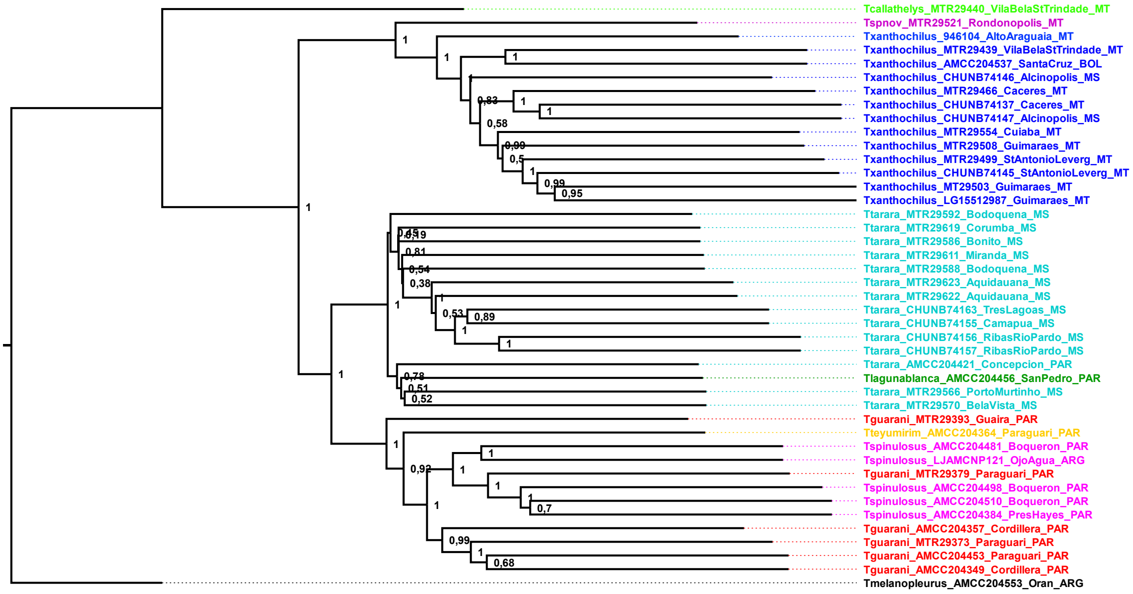


Figure S5. Results produced from the species tree analysis of 1613 UCE loci (matrix with 95% of completeness), produced via ASTRAL. Numbers next to nodes refer to bootstrap support values.

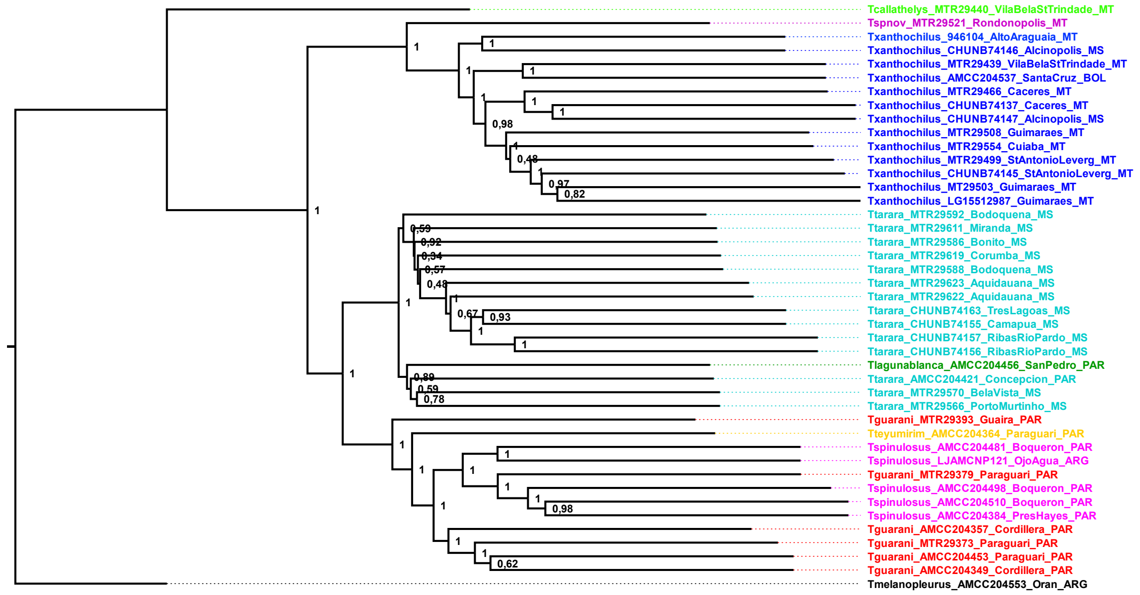


Figure S6. Results produced from the species tree analysis of 2269 UCE loci (matrix with 75% of completeness), produced via ASTRAL. Numbers next to nodes refer to bootstrap support values.

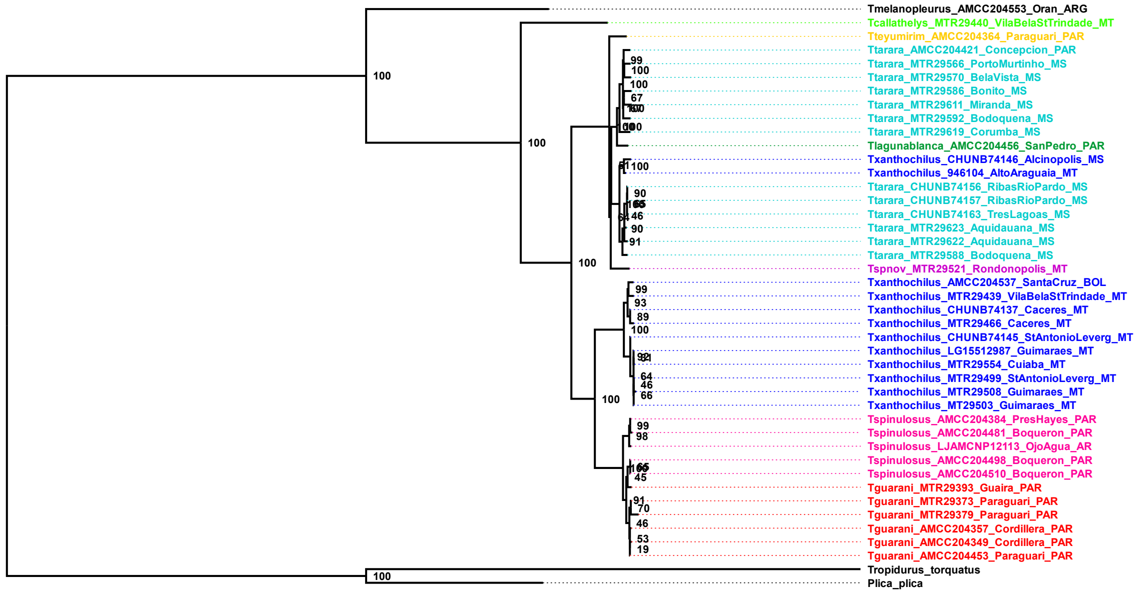


Figure S7. Results produced from the partitioned phylogenetic analysis of 15 mitochondrial loci and 43 samples produced via IQ-TREE. Numbers next to nodes refer to bootstrap support values. The node containing *Plica plica* and *Tropidurus torquatus* was manually set as the outgroup.

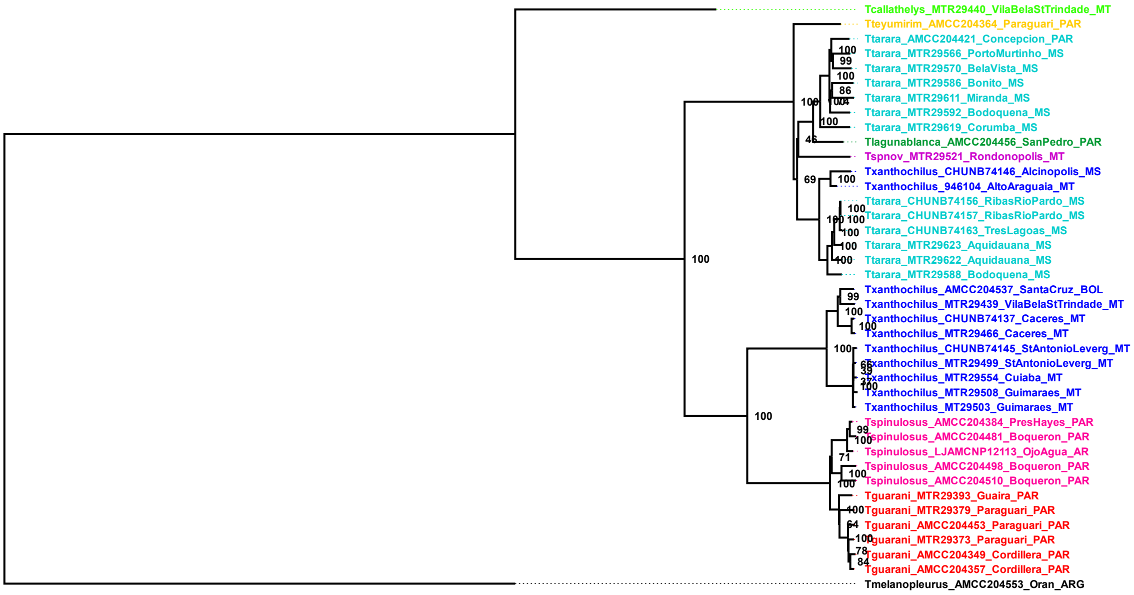


Figure S8. Results produced from the partitioned phylogenetic analysis of 15 mitochondrial loci and 41 samples produced via IQ-TREE, here solely considering species inside the *Tropidurus spinulosus* group (*T. melanopleurus* was manually set as the outgroup). Numbers next to nodes refer to bootstrap support values.

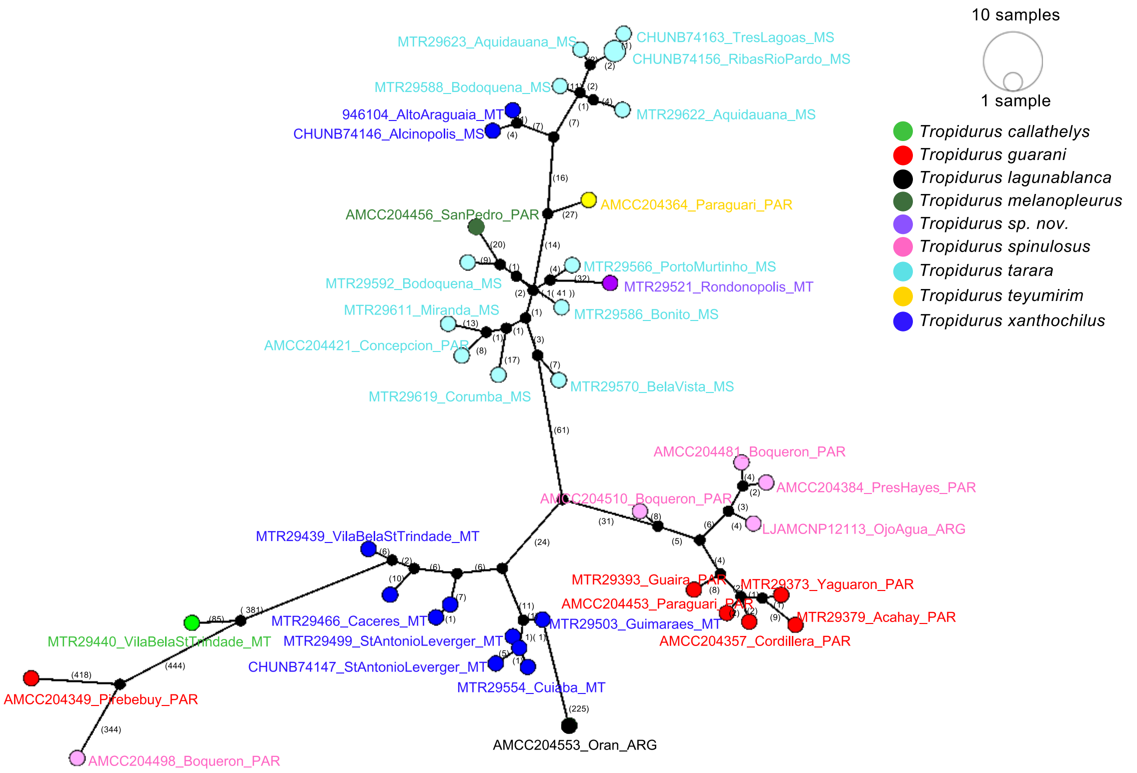


Figure S9. Haplotype network based on three mitochondrial genes (COX1, COX2, and COX3) for 40 individuals belonging to the *Tropidurus spinulosus* species group.

In most cases, the divergence times estimated using RelTime (Figure S11; Tamura et al. 2018) fall within the same 95% highest posterior density (HPD) intervals provided by MCMCTree (dos Reis and Yang 2019), but important differences also occur. For instance, the clade comprising *T. guarani*, *T. spinulosus*, and *T. teyumirim* originates around 4 Ma according to the MCMCTree analysis, while the divergence is estimated to have occurred more recently based on the RelTime approach, around 3 Ma. In contrast, some node ages show notable discrepancies between the two methods. The divergence between *T. sp. nov.* and *T. xanthochilus* is estimated at approximately 3.5 Ma in the MCMCTree analysis, whereas RelTime dates the same split to about 7 Ma. Likewise, the origin of the *T. tarara* clade is estimated at ~3.75 Ma by MCMCTree but nearly 7.5 Ma in the RelTime analysis.

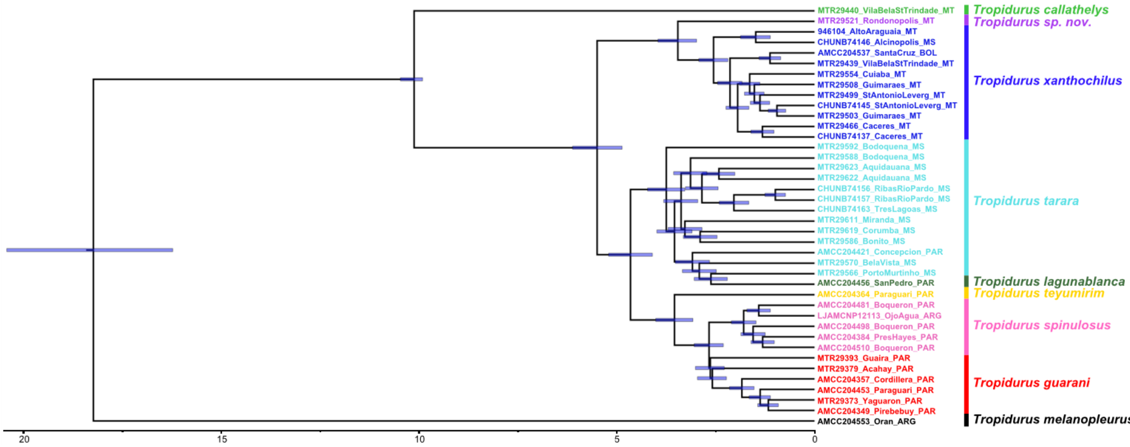

Figure S10. Divergence times (Ma) estimated for the *Tropidurus spinulosus* species group using the UCE dataset. The 95% HPD interval of the posterior estimates (blue shaded bars), as estimated through the MCMCTree program, is shown above each node of the tree. The scale bar is in Ma.

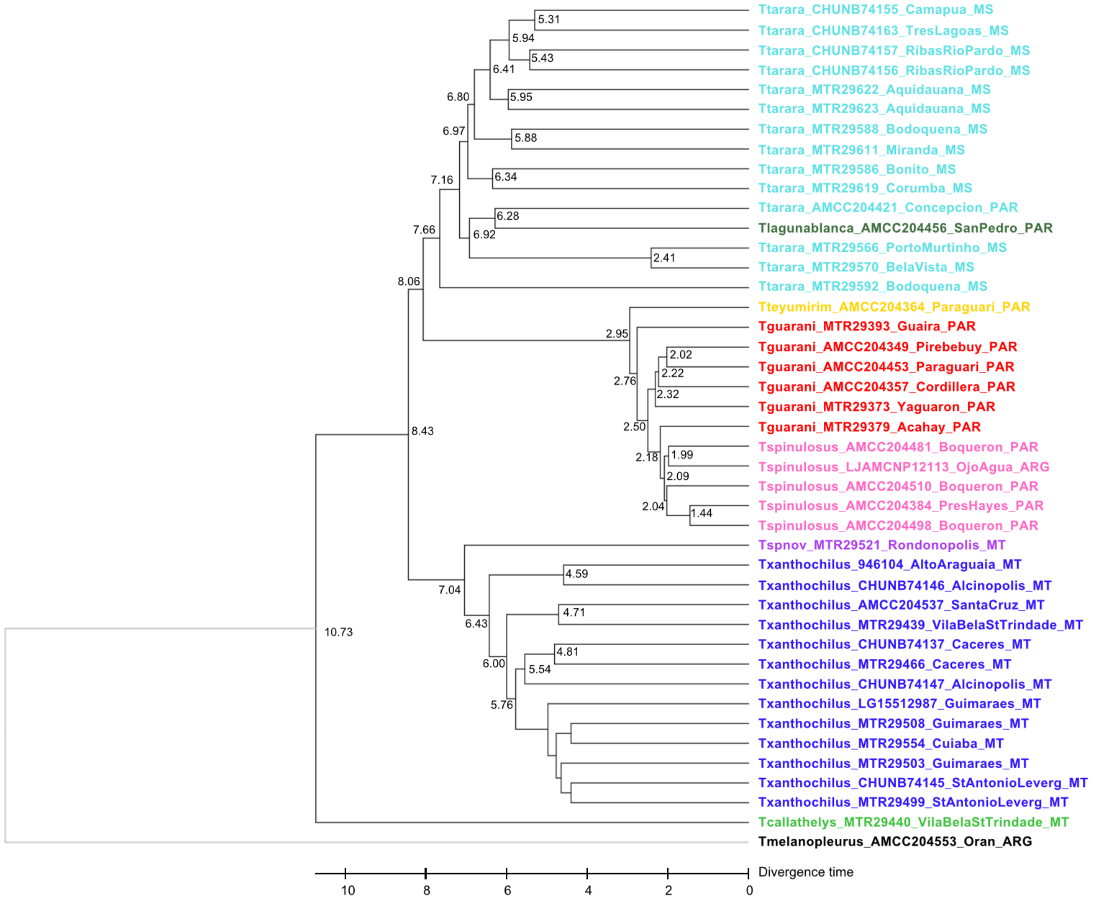


Figure S11. A timetree inferred by applying the RelTime-Branch Length method. The timetree was computed using one calibration constraint, as described in the main text. The scale bar is in Ma.

*Details on phylogenetic networks methodology and results*

Regarding the network approaches, we first used the PhyloNetworks program (Solís-Lemus et al. 2017), in which the inference of phylogenetic networks is done with maximum pseudolikelihood from gene trees or multi-locus sequences (SNaQ; Solís-Lemus and Ané 2016). We supplied SNaQ with unrooted gene trees from 820 UCE loci containing representatives for 32 focal samples; tests were also carried out with 23 and 30 samples, in order to verify the possible influence of the number of tips on the estimation of reticulation events. In all of these alternative tests, we did not use all 43 samples from the UCEs dataset in order to lighten the computational load of the analyses. As results were similar in all configurations, here we only report the estimates with 32 and 30 samples (Figures S13 and S14). Observed concordance factors (CFs) were calculated from the final 820 gene trees dataset (complete matrix, see previous section) and used to infer phylogenetic networks with different maximum numbers of reticulation events (*hmax* up to 12, using five independent runs per h-value) and the proportion (γ) of genomic ancestry in the hybrid lineage contributed by each parental species/lineage. As the default setting for SNaQ considers that each allele in a gene tree corresponds to a specific tip in the network, we also estimated a species-level network, in which each allele/individual was mapped to a particular species, with the assignment done according to the patterns identified through IQ-TREE's partitioned approach. In both cases, we performed a goodness-of-fit test to check whether a given network could explain the quartet concordance factor data adequately (Cai and Ane 2021).

For the PhyloNet analysis (Wen et al. 2018), we applied the MPL algorithm to infer the network, assigning the maximum number of reticulations from 0 to 5, with 10 runs for each reticulation value. We used the '-a taxa map' option to associate taxa with species in the species tree, also following the patterns identified through IQ-TREE's partitioned approach. In this case, all individuals present in the 820 UCE loci dataset were included in the analysis, but using rooted gene trees (using *Tropidurus melanopleurus* as the outgroup). In both PhyloNetworks and PhyloNet analyses, support for each value of *h* (number of hybridization events) was evaluated using the graphical slope heuristic, which involves plotting the pseudolikelihood scores against *h* (referred to as *hmax*), following the framework proposed by Solís-Lemus et al. (2017). Finally, the same dataset used for PhyloNetworks was also used in the NetRAX program (Lutteropp et al. 2022), but the input of NetRAX is the partitioned alignment itself, not the gene trees.

We used D-suite (Malinsky et al. 2021) as a complementary analysis to get gene flow estimates obtained from Patterson's D and f-4 ratio analyses between populations/species. In this approach, ABBA-BABA proportions were obtained with the Dtrios' program for all samples in the *unlinked SNP dataset I,* and applying different assignment configurations of samples to populations/species (Table S10). This program fixes the outgroup and then estimates ABBA-BABA in all possible combinations of three populations. For example, in a tree with four branches plus a fixed outgroup, Dtrios estimates four independent ABBA-BABA by excluding one of the populations at a time, and assigning the remaining three populations as P1, P2, and P3. Moreover, regarding ABBA-BABA positions, in D-suite, P1 and P2 are ordered so that nABBA ≥ nBABA, and the resultant D statistic is always positive. Therefore, all the results, including the f4-ratio and other statistics, reflect evidence of excess allele sharing between P3 and P2 for each trio. Additionally, when testing for introgression among highly divergent species or for ancient introgression events, the D-statistic might indicate false signals of introgression due to substitution rate variation among branches. To address this issue, we used the '--ABBAclustering' option available in the D-suite program, where greater clustering of sites would indicate a real gene flow event rather than a false positive caused by homoplasy.

All phylogenetic network approaches corroborate the pattern of minimal reticulations between species observed in our UCE data, and D-suite tests (Malinsky et al., 2021) for detecting gene flow do not indicate significant nuclear admixture within the *Tropidurus spinulosus* species group as well. First, regarding the PhyloNetworks approach, we obtained a pronounced downward slope in the pseudolikelihood scores (−loglik) as the maximum number of reticulation events (hmax) increased (Figure S12).

More specifically, the relationship between the expected versus observed gene tree discordance among quartets, summarized by concordance factors (CFs), improved when allowing up to eight reticulation events (Figure S13). However, networks with four or more reticulations could not be re-rooted. According to guidelines from PhyloNetworks developers, such networks, despite lower pseudolikelihood values, may not effectively represent the group's phylogeny. Either way, in our network with three reticulations, two occurred between tips of the same species (Figure S14), suggesting that interspecific introgression is not a recurring pattern in the *T. spinulosus* group, at least when it comes to the nuclear genome. Besides, goodness-of-fit tests for all reticulate models showed that all networks failed to provide a good fit to the data, both using the standard option and when allowing multiple alleles per species (Table S1). When permitting multiple alleles, networks with up to five reticulations were produced, with the best fit at $h$ = 1 (Figure S15). However, similar to the results from the standard option, this particular reticulation event also offers little biological insight into potential introgressions within the *T. spinulosus* group.

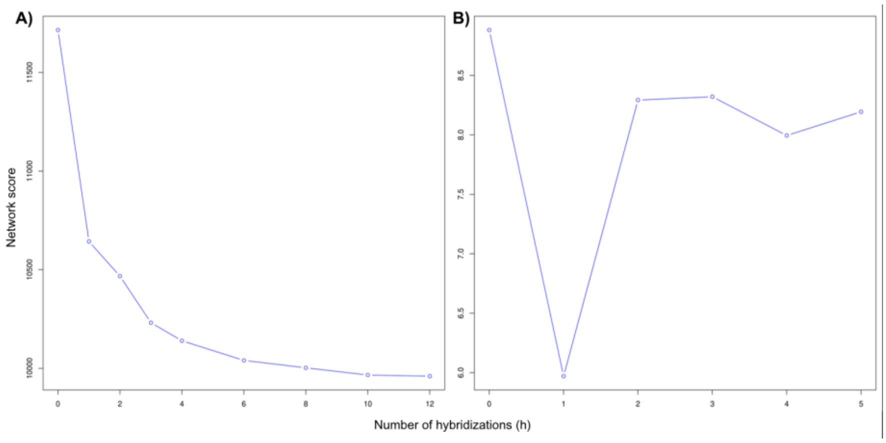


Figure S12. Pseudo-likelihood scores (“network scores”) for each “h” value tested in the PhyloNetworks analysis, using the standard (A) and the multiple alleles option (B). Although better pseudo-likelihood values were obtained for one or more reticulations, it is not recommended to rely on networks that cannot be rooted using your outgroup clade. These networks are likely incorrect because, based on external information, the indicated outgroup is indeed an outgroup. In our case, this issue applies to all scenarios beyond three reticulations. Furthermore, even allowing more than eight reticulations (up to 12), networks with a maximum of 8 reticulations were obtained.

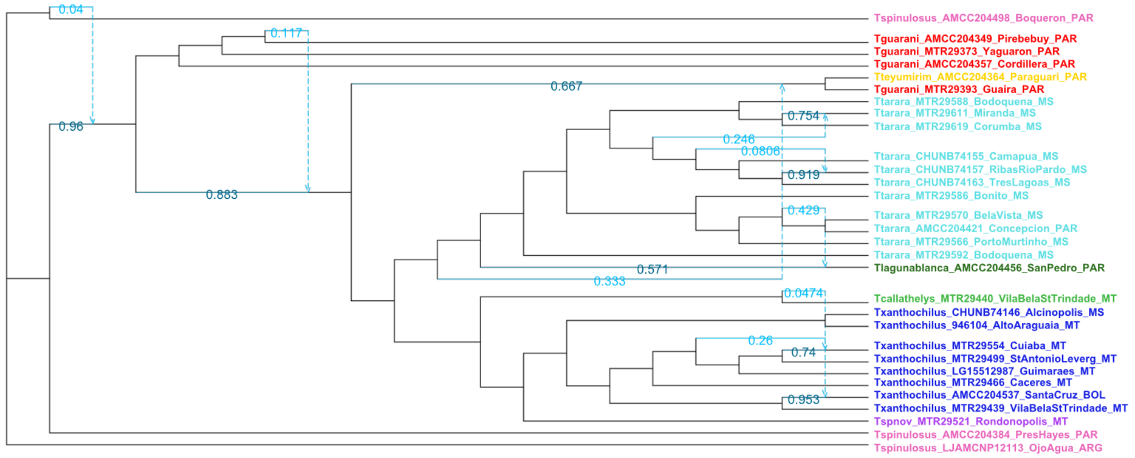


Figure S13. Unrooted phylogenetic network of the *Tropidurus spinulosus* species group (30 samples) produced through the PhyloNetworks program, in the standard option with eight potential reticulations. Blue branches (both solid and dashed) indicate inferred hybridization events, and numbers next to the branches show the estimated proportion of genes contributed by each lineage in the hybridization event. It is important to note that even if it were possible to trust this network (which is not the case, as re-rooting was not possible), practically no introgression information would be added, as most reticulations occur within species.

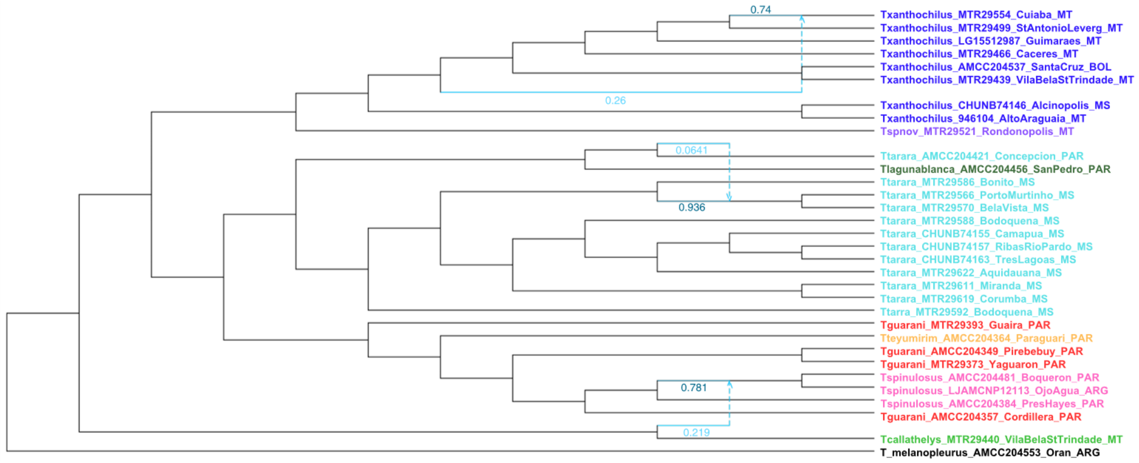


Figure S14. Phylogenetic network of *Tropidurus spinulosus* species group (UCE dataset, 32 samples) with three potential reticulations (hmax=3). Blue branches (both solid and dashed) indicate inferred hybridization events, and numbers next to the branches show the estimated proportion of genes contributed by each lineage in the introgression event.

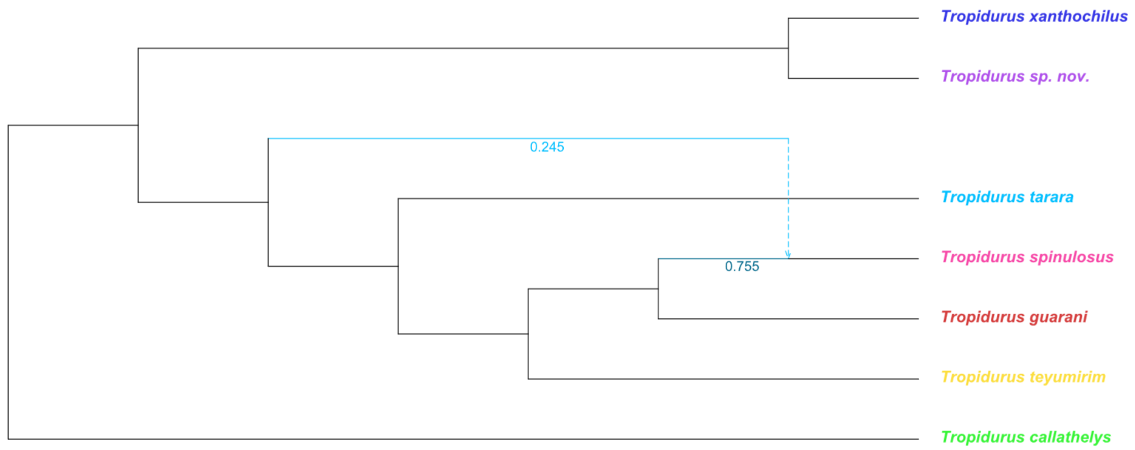


Fig. S15. Unrooted phylogenetic network of the *Tropidurus spinulosus* species group (30 samples) produced through the PhyloNetworks program, in the multiple alleles option with a single reticulation. Blue branches (both solid and dashed) indicate inferred hybridization events, and numbers next to the branches show the estimated proportion of genes contributed by each lineage in the hybridization event. It is important to note that even if it were possible to trust this network (which is not the case, as re-rooting was not possible), practically no introgression information would be added, as most reticulations occur within species.

Table S1. Statistical values associated with the goodness-of-fit tests applied to phylogenetic network models (here represented as the number of reticulations, or *h* values) analyzed with the PhyloNetworks program. In this test, more likely network topologies are expected to have higher values (at least >0.05), corresponding to a high proportion of well-fit quartet trees.

| SNaQ! option | *h* = 0 | 1 | 2 | 3 | 4 | 5 | 6 | 8 |
|---|---|---|---|---|---|---|---|---|
| Standard | 1.9e-216 | 2.5e-182 | 9.35e-201 | 9.8e-200 | 5.9e-140 | - | 1.0e-97 | 3.6e-114 |
| Multiple alleles | 6.86e-5 | 5.42e-2 | 1.099e-3 | 9.89e-4 | 8.44e-4 | 9.79e-4 | - | - |

Employing the PhyloNet approach (Wen et al. 2018), the species tree with the best pseudolikelihood value (namely, the lower value) was one without any reticulations (Figures S16 and S17). Using NetRax (Lutteropp et al. 2022), the optimal network inferred included five reticulations. However, again, the majority of these reticulations occurred between tips of the same species, with only one reticulation detected between different species (Figure S18). Given that NetRax does not account for ILS, this interspecific reticulation may be an artifact of this limitation.

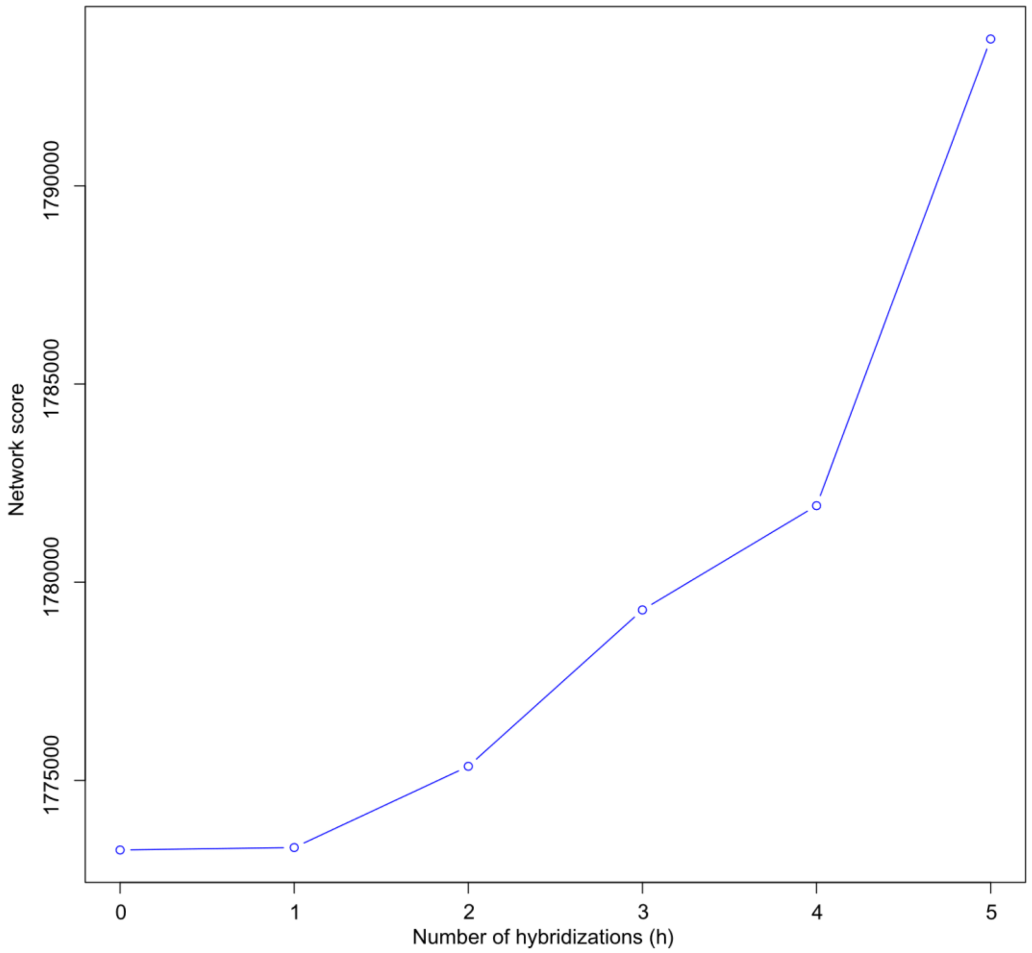


Figure S16. Pseudo-likelihood scores (here as “network scores”) for each h value tested in the PhyloNet analysis.

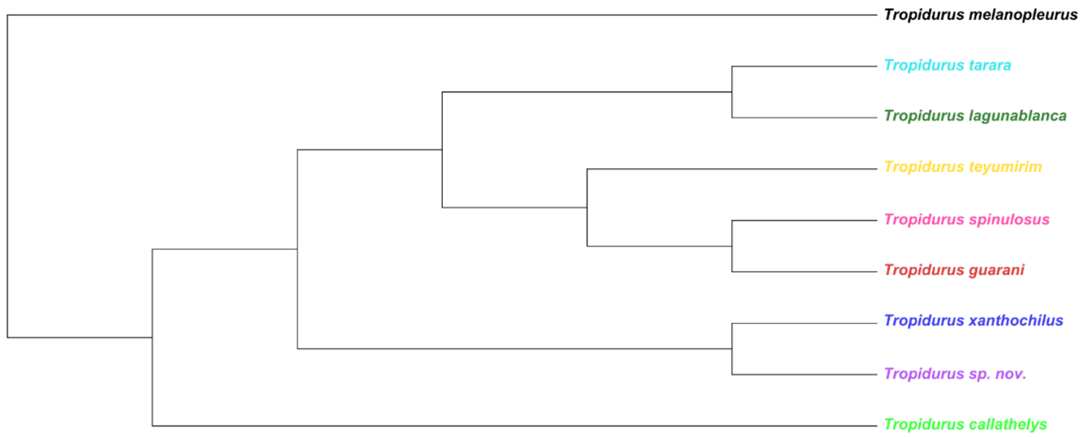


Figure S17. Optimal phylogenetic network of the *Tropidurus spinulosus* species group (UCE dataset) produced through the PhyloNet program, in this case with no reticulations.

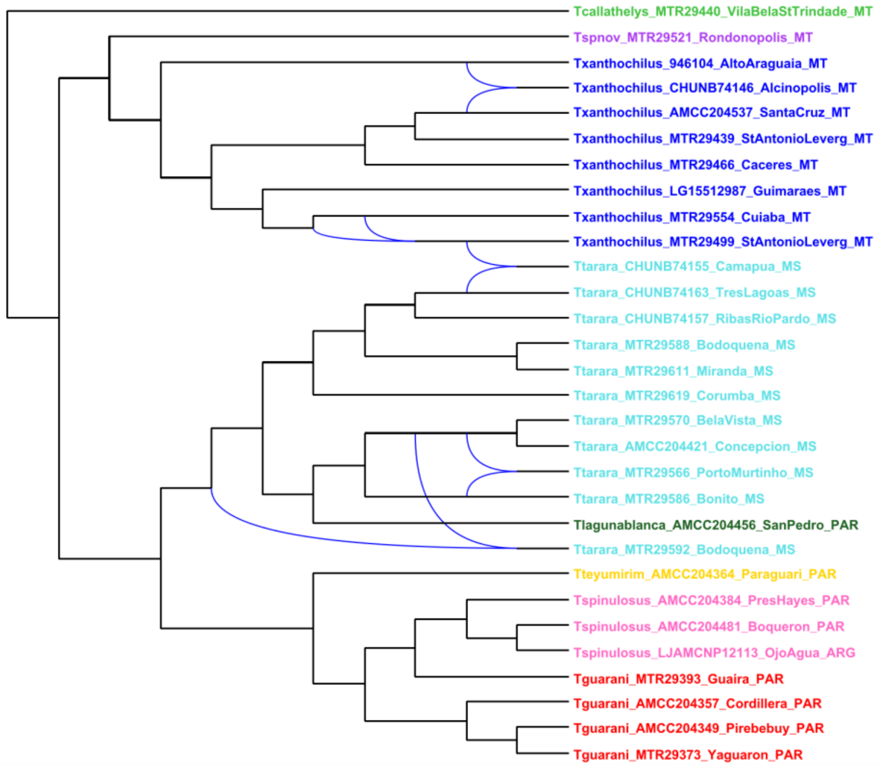


Figure S18. Phylogenetic network of the *Tropidurus spinulosus* species group produced through the NetRax program. In this case, there are five potential reticulation events (shown by blue links between branches), but only one occurs between tips of different species.

Consistent with the phylogenetic network's analyses, D-suite tests (Malinsky et al., 2021) for detecting gene flow do not indicate significant nuclear admixture within the *Tropidurus spinulosus* species group. Preliminary D-statistic calculations suggested possible introgression; however, after accounting for variation in substitution rates across phylogenetic branches using the "sensitive" test available in the '--ABBAclustering' option of D-suite, only one trio remained significant (*p*-value < 0.05), indicating gene flow between *T. teyumirim* and *T. tarara* (Table S2).

Table S2. ABBA-BABA test values and related statistics used to assess evidence of gene flow between populations or closely related species (here referred to as P1, P2, and P3). Only significant results (*p value* < 0.05) from the primary analysis and the ABBA-clustering test are shown (reflecting an excess allele sharing between P2 and P3 for each trio). For more details surrounding each assignment category, please check the Supplementary Material.

| P1 | P2 | P3 | D statistic | f4-ratio | BBAA | ABBA | BABA | Assignment category |
|---|---|---|---|---|---|---|---|---|
| guaranigr* | teyumirim | tararagr | 0.287931 | 0.210312 | 134.129 | 50.685 | 28.022 | 5spp |
| guarani | teyumirim | tararagr | 0.18045 | 0.143375 | 136.144 | 46.461 | 32.257 | 6spp |
| guarani | teyumirim | tararagr | 0.219168 | 0.151037 | 144.779 | 44.692 | 28.623 | 8spp |

* The abbreviation 'gr' used here does not necessarily denote a monophyletic group. It is simply a way to indicate that, in this case, *T. teyumirim* was considered within the same grouping as the specific samples of *T. guarani*. The same applies to *'tararagr'*, which includes the specific samples of *T. tarara* as well as the *T. lagunablanca* sample.

*Migration tests in BPP*

Our BPP population size estimates reveal significant differences among ancestral populations likely involved in the ancient introgression events underlying the observed discordance. These differences also support the inferred directionality of gene flow. For example, the highest estimated interspecific migration rate when using the mitochondrial topology as the guide tree was M(*T. xanthochilus* → ancestor of *T. guarani* + *T. spinulosus*) = 0.3629 (Figure 3, Table S3). In this case, the θ estimate of *T. xanthochilus* was considerably lower than that of the ancestral population of *T. guarani* and *T. spinulosus* (θ = 0.0006 and 0.2017, respectively; Table S5), further supporting the plausibility of mitochondrial capture event—where some *T. xanthochilus* mitogenomes might have been assimilated by the larger emerging *T. guarani* + *T. spinulosus* clade. Similarly, when using nuclear topology as a guide tree, the largest migration rate involving *T. xanthochilus* was with the ancestor of the ((*T. guarani*, *T. spinulosus*), *T. tarara*)) clade (Figure 3, Table S4). Accordingly, the estimated θ value was larger in the ancestral node compared to *T. xanthochilus* (θ = 0.0045 and 0.0020, respectively; Table S6), once again suggesting an ancient introgression between these lineages.

Table S3. BPP migration rates estimates, ordered from highest to lowest values, obtained using the mitochondrial topology as guide tree and the nuclear dataset as input. Ancestral nodes were categorized as follows: A: (*Tropidurus xanthochilus* 1, *T. tarara* 2) ancestral; B: (*T. xanthochilus* 2, (*T. spinulosus*, *T. guarani*) ancestral; C: (*T. guarani*, *T. spinulosus*) ancestral;

| Migration rates | Migration rate | Standard deviation |
|---|---|---|
| *T. xanthochilus* 1 → *T. xanthochilus* 2 | 0.0229 | 0.0176 |
| *T. xanthochilus* 2 → *T. xanthochilus* 1 | 0.3732 | 0.0602 |
| *T. xanthochilus* 1 → C | 0.3629 | 0.2653 |
| C → *T. xanthochilus* 1 | 0.1810 | 0.087 |
| A → C | 0.0999 | 0.1000 |
| C → A | 0.1001 | 0.1000 |
| *T. xanthochilus* 2 → A | 0.0917 | 0.0917 |

| | | |
|---|---|---|
| A → *T. xanthochilus* 2 | 0.0546 | 0.0624 |
| A → B | 0.0778 | 0.0787 |
| B → A | 0.0956 | 0.0955 |
| *T. xanthochilus* 1 → B | 0.0188 | 0.0202 |
| B → *T. xanthochilus* 1 | 0.0222 | 0.0238 |
| *T. xanthochilus* 2 → B | 0.0073 | 0.0073 |
| B → *T. xanthochilus* 2 | 0.0064 | 0.0066 |
| *T. xanthochilus* 1 → *T. tarara* 2 | 0.0053 | 0.0048 |
| *T. tarara* 2 → *T. xanthochilus* 1 | 0.0031 | 0.0031 |
| *T. xanthochilus* 1 → *T. guarani* | 0.0037 | 0.0037 |
| *T. guarani* → *T. xanthochilus* 1 | 0.0032 | 0.0032 |
| *T. spinulosus* → *T. xanthochilus* 2 | 0.0028 | 0.0024 |
| *T. xanthochilus* 2 → *T. spinulosus* | 0.0022 | 0.0022 |
| *T. xanthochilus* 1 → *T. spinulosus* | 0.0024 | 0.0024 |
| *T. spinulosus* → *T. xanthochilus* 1 | 0.0033 | 0.0033 |
| *T. guarani* → *T. xanthochilus* 2 | 0.0018 | 0.0018 |
| *T. xanthochilus* 2 → *T. guarani* | 0.0028 | 0.0028 |

Table S4. BPP migration rates estimates, ordered from highest to lowest values, using the nuclear topology as the guide tree and incorporating both nuclear and mitochondrial datasets as input. Ancestral nodes were categorized as follows: B: (*T. guarani*, *T. spinulosus*) ancestral; D: (*T. xanthochilus*, *T. sp nov.*) ancestral; E: (*T. tarara*, (*T. teyumirim*, (*T. guarani*, *T. spinulosus*))) ancestral; F: (*T. teyumirim*, (*T. guarani*, *T. spinulosus*)) ancestral. Values shown as '–' indicate parameters that remained fixed during the BPP run.

| Migration rates | nuDNA only | | nuDNA + mtDNA | |
|---|---|---|---|---|
| | Migration rate \| Standard deviation | | | |
| *T. xanthochilus* → E | 0.0813 | 0.0752 | 0.3030 | 0.1467 |
| E → *T. xanthochilus* | 0.0824 | 0.0824 | 0.0656 | 0.0664 |
| E → D | 0.0346 | 0.0312 | 0.2400 | 0.2377 |
| D → E | 0.0261 | 0.0240 | 0.0289 | 0.0233 |
| D → B | 0.0951 | 0.0886 | - | - |
| B → D | 0.0922 | 0.0854 | - | - |
| *T. tarara* → D | 0.0652 | 0.0711 | - | - |
| D → *T. tarara* | 0.0796 | 0.0742 | - | - |
| D → F | 0.0501 | 0.0637 | - | - |
| F → D | 0.0592 | 0.0641 | - | - |
| *T. xanthochilus* → F | 0.0175 | 0.0271 | 0.0276 | 0.0162 |
| F → *T. xanthochilus* | 0.0320 | 0.0369 | 0.0347 | 0.0243 |
| *T. xanthochilus* → B | 0.0175 | 0.0238 | 0.0153 | 0.0181 |
| B → *T. xanthochilus* | 0.0157 | 0.0172 | 0.0113 | 0.0120 |

| | | | | |
|---|---|---|---|---|
| *T. xanthochilus* → *T. tarara* | 0.0087 | 0.0065 | 0.0116 | 0.0065 |
| *T. tarara* → *T. xanthochilus* | 0.0060 | 0.0042 | 0.0097 | 0.0048 |
| *T. xanthochilus* → *T. spinulosus* | 0.0026 | 0.0025 | 0.0025 | 0.0024 |
| *T. spinulosus* → *T. xanthochilus* | 0.0040 | 0.0036 | 0.0035 | 0.0032 |
| *T. xanthochilus* → *T. guarani* | 0.0027 | 0.0027 | 0.0026 | 0.0026 |
| *T. guarani* → *T. xanthochilus* | 0.0030 | 0.0029 | 0.0042 | 0.0037 |
| *T. spinulosus* → D | - | - | - | - |
| D → *T. spinulosus* | - | - | - | - |
| *T. guarani* → D | - | - | - | - |
| D → *T. guarani* | - | - | - | - |

Table S5. Population sizes estimated through BPP for species and nodes involved in the discordance, obtained using the mitochondrial topology as a guide tree and the nuclear dataset as input. Ancestral nodes were categorized as follows: A: (*Tropidurus xanthochilus* 1, *T. tarara* 2) ancestral; B: (*T. guarani*, *T. spinulosus*) ancestral; C: (*T. xanthochilus* 2, (*T. spinulosus*, *T. guarani*)) ancestral.

| Population/species | Θ estimate | Standard deviation |
|---|---|---|
| A | 3.6850e-2 | 3.4800e-2 |
| C | 2.0170e-2 | 1.7100e-2 |
| *Tropidurus tarara* 1 | 2.8692e-3 | 2.7064e-4 |
| B | 2.1970e-3 | 2.4250e-4 |
| *Tropidurus tarara* 2 | 2.1579e-3 | 1.6417e-4 |
| *Tropidurus guarani* | 1.8066e-3 | 1.4787e-4 |
| *Tropidurus xanthochilus* 2 | 1.3980e-3 | 7.9320e-5 |
| *Tropidurus spinulosus* | 1.0771e-3 | 8.7154e-5 |
| *Tropidurus callathelys* | 9.9579e-4 | 1.2489e-4 |
| *Tropidurus teyumirim* | 8.4267e-4 | 9.8651e-5 |
| *Tropidurus melanopleurus* | 8.3996e-4 | 9.9519e-5 |
| *Tropidurus sp. nov.* | 6.8688e-4 | 7.8393e-5 |
| *Tropidurus xanthochilus* 1 | 6.2580e-4 | 6.0670e-5 |

Table S6. Population sizes estimated through BPP for species and nodes involved in the discordance, obtained using the nuclear topology as guide tree, and both the nuclear and mitochondrial datasets as input. Ancestral nodes were categorized as follows: B: (*T. guarani*, *T. spinulosus*) ancestral; D: (*T. xanthochilus*, *T. sp nov.*) ancestral; E: (*T. tarara*, (*T. teyumirim*, (*T. guarani*, *T. spinulosus*))) ancestral; F: (*T. teyumirim*, (*T. guarani*, *T. spinulosus*)) ancestral.

| Population/species | nuDNA only | | nuDNA + mtDNA | |
|---|---|---|---|---|
| | Θ estimate | Standard deviation | | |
| B | 5.5213e-3 | 3.3508e-3 | 6.2000e-3 | 2.6615e-3 |
| *Tropidurus tarara* | 5.1255e-3 | 2.3624e-4 | 5.1733e-3 | 2.4583e-4 |
| E | 4.4514e-3 | 5.4385e-4 | 4.8863e-3 | 6.2706e-4 |
| D | 2.5391e-3 | 3.3727e-4 | 3.1248e-3 | 5.4175e-4 |
| *Tropidurus xanthochilus* | 2.01480-3 | 1.1679e-4 | 2.1753e-3 | 1.2265e-4 |
| F | 2.0132e-3 | 3.0304e-4 | 5.2742e-4 | 4.7758e-5 |
| *Tropidurus guarani* | 1.6841e-3 | 1.448e-4 | 1.6183e-3 | 1.4108e-4 |
| *Tropidurus spinulosus* | 1.024e-3 | 8.5984e-5 | 1.0046e-3 | 8.4714e-5 |
| *Tropidurus callathelys* | 9.9434e-4 | 1.2486e-4 | 9.8606e-4 | 1.2319e-4 |
| *Tropidurus melanopleurus* | 8.4099-4 | 9.9533e-5 | 8.2760e-4 | 9.7530e-5 |
| *Tropidurus sp. nov.* | 8.3069e-4 | 9.9668e-5 | 8.4784e-4 | 1.0065e-4 |
| *Tropidurus teyumirim* | 8.0998e-4 | 9.9482e-5 | 7.9985e-4 | 9.7108e-5 |

*Details on demographic inference methodology and results*

To estimate changes in effective population sizes over time using the mitochondrial dataset, we performed the multilocus coalescent-based Extended Bayesian Skyline Plots (EBSP; Heled and Drummond 2008) implemented in BEAST v2.7.6 (Bouckaert et al. 2019). Four lineages were used in this analysis, namely: *Tropidurus guarani*, *T. spinulosus*, one *T. tarara* clade, and *T. xanthochilus*. Regarding this, as mentioned in the main text, we highlight that demographic analyses typically assume that each analyzed group is monophyletic and below the species level. While *T. guarani* is consistently recovered as paraphyletic in our nuclear phylogenies, it forms a monophyletic clade sister to *T. spinulosus* in mitochondrial analyses. Therefore, for the

EBSP analysis, which was performed exclusively with mitochondrial data, *T. guarani* was treated as a single lineage. A similar issue arises with *T. tarara*, which appears as two distinct mitochondrial groups. In this case, analyses were conducted separately for each cluster, but since the results were identical, only one is presented in the text, as it effectively represents both *T. tarara* groupings. In the remaining cases, samples were divided according to the clades recovered in the mtDNA partitioned approaches previously described.

To reduce computational burden and facilitate convergence, we conducted separate analyses on three sets of mitochondrial genes rather than using the entire mitochondrial genome. These sets included: 1) cytochrome b (cytb); 2) COX1, COX2, and COX3; 3) 12S and 16S ribosomal RNAs. For coding gene analyses, alignments were partitioned by gene and codon position, with tree and clock models linked across all partitions. Site models were treated independently, with BEAST Model Test performed separately for each partition to estimate mutation rates, and restricting potential models to the "namedExtended" set. Regarding substitution rates, we set $1.0 \times 10^{-2}$ as the mean for the COX partition, $6.339 \times 10^{-3}$ for the 12s + 16s partition, and $1.94 \times 10^{-2}$ for the cytb partition, all reference values already used in previous studies that used similar approaches when dealing with *Tropidurus’* mtDNA in BEAST (e.g., Werneck et al. 2015; Camurugi et al. 2022; Carvalho et al. 2024). An uncorrelated relaxed molecular clock was adopted in all three cases (with a normal distribution for the ‘ucldMean’; Sigma = 0.0025). Because the default of the hyper-prior of the mean of the population size distribution ("populationMean.alltrees") is a sensitive option and takes a long time to converge (requiring longer runs), we modified the prior to a Normal distribution centered on 1.0 (Mean) with a standard deviation of 0.1 (Sigma) to improve convergence. Initial tests were carried out to determine the optimum number of MCMC simulations required for each of the populations. In most tests, we ran 500,000,000 MCMC simulations, sampled every 50,000 chains. In all cases, three independent runs were carried out, and we checked for convergence using Tracer 1.7 (Rambaut et al. 2018) by visually checking effective sample size (ESS > 200). In specific cases where convergence was not achieved with this number of simulations, we combined logs with LogCombiner (Rambaut and Drummond 2014) and used those files to generate EBSP plots.

To infer the demographic history using the nuDNA dataset, we first generated a folded SFS file for the SNP dataset with easySFS (Gutenkunst et al. 2009). We then used Stairway Plot v. 2.18 (Liu and Fu 2020) to estimate historical effective population size changes of *Tropidurus xanthochilus*, *T. tarara*, *T. spinulosus,* and *T. guarani*. For this last group, only the four samples that form a clade in the partitioned phylogenetic analyses described above were used (Figures S1-S6). The parameters for the Stairway Plot analysis were as follows: 67% for the training sites, and one year was assumed as generation time (A. Carvalho, pers. commun.). The nuclear mutation rate was assumed to be $7.17 \times 10^{-9}$ substitutions/site/generation, which is the value available for the phylogenetically closest species to *Tropidurus* in the germline mutation rate study of Bergeron et al. (2023).

Our EBSPs revealed a pronounced increase in effective population size for all populations, particularly in the last 100.000 years (Figure S19). Given that *Tropidurus tarara* was recovered in two distinct clusters within the mitochondrial tree, we reinforce that independent analyses were conducted for each group. As results were identical across both groups, we present the result for only one cluster, which we consider representative of the species as a whole. For the nuclear dataset, Stairway Plot analyses for *T. xanthochilus*, *T. tarara*, *T. spinulosus*, and *T. guarani* revealed a consistent pattern across species, characterized by population expansion mainly beginning 60,000 years ago, followed by relatively stable population sizes thereafter (Figure S20).

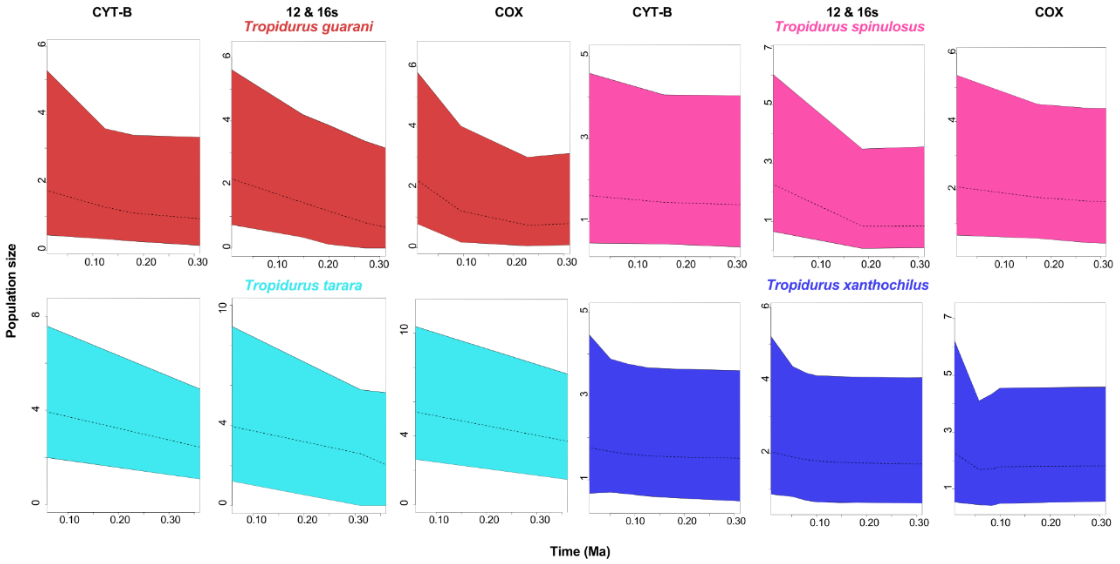


Figure S19. EBSP results showing changes in effective population size through time for lineages in the *Tropidurus spinulosus* species group, based on three mitochondrial partitions (from left to right: cytb, 12 + 16s, and COX1, 2, and 3). Y-axis shows the effective population size with respective mean (dashed lines) and 95% HPD limits (the surrounding colored area delimited by the continuous lines). Assuming a generation time of 1 year, the current effective population size estimated at one would be equivalent to around 1 million. While this number appears large, it serves only as an illustration. Actual values will vary depending on the chosen mutation rate and generation time. The x-axis shows time in Ma as time goes backwards from left to right.

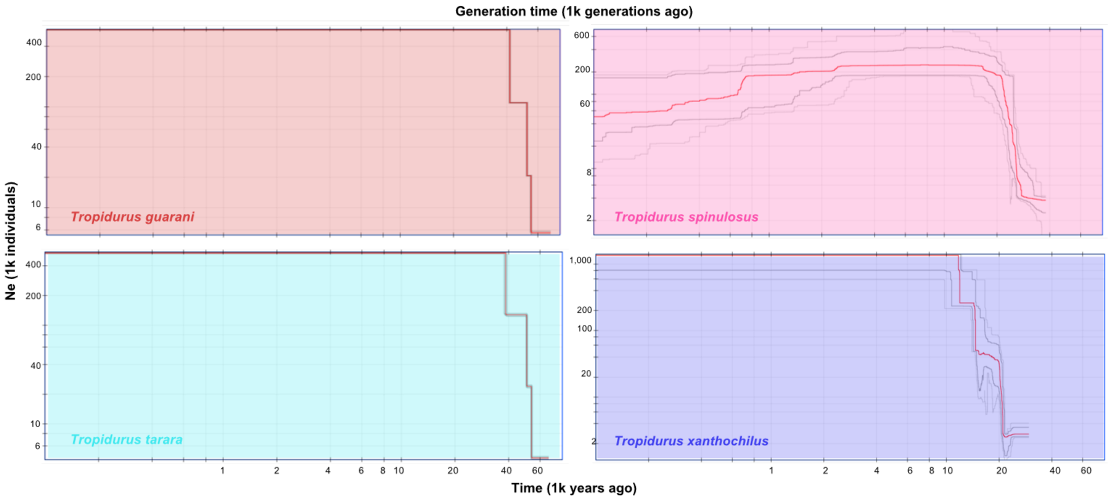


Figure S20. Stairway Plot demographic inferences of four lineages within the *Tropidurus spinulosus* species group. The red line shows the median estimate of the effective population size (Ne). The gray lines indicate the 75% (dark) and 95% (light) confidence intervals.

### *Remaining supplementary tables*

Table S7. Nucleotide diversity values (π) for species within the *Tropidurus spinulosus* group, based on their mitogenomes.

| Population/species | Average number of nucleotide differences per site |
|---|---|
| *Tropidurus guarani* | 0.005966946 |
| *Tropidurus spinulosus* | 0.007992678 |
| *Tropidurus tarara* | 0.00638382 |
| *Tropidurus xanthochilus* | 0.009012845 |

Table S8. Tajima's D results.

| Population/species | D value | p-value |
|---|---|---|
| *Tropidurus guarani* | -5.481613 | 4.214645 x 10-8 |
| *Tropidurus spinulosus* | -7.461749 | 8.538173 x 10-14 |
| *Tropidurus tarara* | -3.737147 | 1.861204 x 10-4 |
| *Tropidurus xanthochilus* | -4.729045 | 2.255783 x 10-6 |

Table S9. Sampling localities and geographical coordinates for the 43 specimens included in this study, alongside their genetic assignment in the BEAST analysis.

| Sample ID | General species assignment | BEAST assignment | Location | Longitude | Latitude |
|---|---|---|---|---|---|
| MTR 29623 | *Tropidurus tarara* | tarara2 | Aquidauana, Mato Grosso do Sul, BRA | -55.5543 | -20.4830 |
| MTR 29622 | *Tropidurus tarara* | tarara2 | Aquidauana, Mato Grosso do Sul, BRA | -55.8217 | -20.4445 |
| MTR 29619 | *Tropidurus tarara* | tarara | Corumbá, Mato Grosso do Sul, BRA | -57.0361 | -19.5738 |
| MTR 29611 | *Tropidurus tarara* | tarara | Miranda, Mato Grosso do Sul, BRA | -56.5084 | -20.1928 |
| MTR 29592 | *Tropidurus tarara* | tarara | Bodoquena, Mato Grosso do Sul, BRA | -56.8965 | -20.5319 |
| MTR 29588 | *Tropidurus tarara* | tarara | Bodoquena, Mato Grosso do Sul, BRA | -56.7138 | -20.5424 |
| MTR 29586 | *Tropidurus tarara* | tarara | Bonito, Mato Grosso do Sul, BRA | -56.5812 | -21.1074 |
| MTR 29570 | *Tropidurus tarara* | tarara | Bela Vista, Mato Grosso do Sul, BRA | -56.4635 | -21.9755 |
| MTR 29566 | *Tropidurus tarara* | tarara | Porto Murtinho, Mato Grosso do Sul, BRA | -57.8963 | -21.7065 |
| MTR 29554 | *Tropidurus xanthochilus* | xanthochilus | Cuiabá, Mato Grosso, BRA | -55.9886 | -15.4575 |
| MTR 29521 | *Tropidurus sp. nov.* | spnov | Rondonópolis, Mato Grosso, BRA | -54.7676 | -16.6510 |
| MTR 29508 | *Tropidurus xanthochilus* | xanthochilus | Chapada dos Guimarães, Mato Grosso, BRA | -55.8313 | -15.4072 |
| MTR 29503 | *Tropidurus xanthochilus* | xanthochilus | Chapada dos Guimarães, Mato Grosso, BRA | -55.7723 | -15.4490 |

| | | | | | |
|---|---|---|---|---|---|
| MTR 29499 | *Tropidurus xanthochilus* | xanthochilus | Santo Ant. do Leverger, Mato Grosso, BRA | -55.5380 | -15.9643 |
| MTR 29466 | *Tropidurus xanthochilus* | xanthochilus | Cáceres, Mato Grosso, BRA | -57.2507 | -15.6263 |
| MTR 29440 | *Tropidurus callathelys* | callathelys | Vila Bela da St. Trindade, Mato Grosso, BRA | -60.0703 | -14.9111 |
| MTR 29439 | *Tropidurus xanthochilus* | xanthochilus | Vila Bela da St. Trindade, Mato Grosso, BRA | -59.9546 | -15.0086 |
| MTR 29393 | *Tropidurus guarani* | guarani | Villarrica, Guairá, PAR | -56.2890 | -25.8323 |
| MTR 29379 | *Tropidurus guarani* | guarani | Acahay, Paraguarí, PAR | -57.1733 | -25.8942 |
| MTR 29373 | *Tropidurus guarani* | guarani | Yaguarón, Paraguarí, PAR | -57.2941 | -25.5675 |
| LJAM-CNP 12113 | *Tropidurus spinulosus* | spinulosus | Ojo de Água, ARG | -63.6961 | -29.5004 |
| LG1551/2987 | *Tropidurus xanthochilus* | spinulosus | Chapada dos Guimarães, Mato Grosso, BRA | -55.8000 | -14.8666 |
| CHUNB74163 | *Tropidurus tarara* | tarara2 | Três Lagoas, Mato Grosso do Sul, BRA | -52.1941 | -20.5516 |
| CHUNB74157 | *Tropidurus tarara* | tarara2 | Ribas Rio Pardo, Mato Grosso do Sul, BRA | -53.4730 | -20.2532 |
| CHUNB74156 | *Tropidurus tarara* | tarara2 | Ribas Rio Pardo, Mato Grosso do Sul, BRA | -53.4730 | -20.2532 |
| CHUNB74155 | *Tropidurus tarara* | - | Camapuã, Mato Grosso do Sul, BRA | -54.1364 | -19.3246 |
| CHUNB74147 | *Tropidurus xanthochilus* | - | Alcinópolis, Mato Grosso do Sul, BRA | -53.4039 | -18.8577 |
| CHUNB74146 | *Tropidurus xanthochilus* | - | Alcinópolis, Mato Grosso do Sul, BRA | -53.4325 | -18.6444 |
| CHUNB74145 | *Tropidurus xanthochilus* | xanthochilus | Santo Ant. do Leverger, Mato Grosso, BRA | -55.3031 | -15.5231 |
| CHUNB74137 | *Tropidurus xanthochilus* | xanthochilus | Cáceres, Mato Grosso, BRA | -57.3437 | -16.3297 |
| AMCC 204553 | *Tropidurus melanopleurus* | melanopleurus | Águas Blancas, Oran, ARG | -64.3696 | -22.7274 |
| AMCC 204537 | *Tropidurus xanthochilus* | xanthochilus | Santa Cruz, BOL | -61.0344 | -14.7665 |
| AMCC 204510 | *Tropidurus spinulosus* | spinulosus | Boquerón, PAR | -60.5332 | -20.3763 |
| AMCC 204498 | *Tropidurus spinulosus* | spinulosus | Boquerón, PAR | -61.6625 | -21.2040 |
| AMCC 204481 | *Tropidurus spinulosus* | spinulosus | Loma Plata, Boquerón, PAR | -59.8611 | -22.3435 |
| AMCC 204456 | *Tropidurus lagunablanca* | tarara | Santa Rosa del Araguay, San Pedro, PAR | -56.2946 | -23.8120 |
| AMCC 204453 | *Tropidurus guarani* | guarani | Paraguarí, PAR | -57.1296 | -25.6068 |
| AMCC 204421 | *Tropidurus tarara* | tarara | Concepcíon, PAR | -57.3693 | -22.6923 |
| AMCC 204384 | *Tropidurus spinulosus* | spinulosus | Rio Verde, Presidente Hayes, PAR | -59.2027 | -23.2140 |
| AMCC 204364 | *Tropidurus teyumirim* | teyumirim | Paraguarí, PAR | -56.8702 | -26.0499 |

| | | | | | |
|---|---|---|---|---|---|
| AMCC 204357 | *Tropidurus guarani* | guarani | San Bernardino, Cordillera, PAR | -57.3027 | -25.3071 |
| AMCC 204349 | *Tropidurus guarani* | guarani | Pirebebuy, Cordillera, PAR | -57.0456 | -25.5144 |
| 946104 | *Tropidurus xanthochilus* | - | Alto Araguaia, Mato Grosso, BRA | -53.2207 | -17.3136 |

Table S10. Different assignment configurations of samples to populations/species for all D-suite analyses conducted on this study.

| Sample | Assigned population 1 | Assigned population 2 | Assigned population 3 | Assigned population 4 |
|---|---|---|---|---|
| AMCC 204456 | tararagr | tararagr | tararagr | tararagr |
| MTR 29570 | tararagr | tararagr | tararagr | tararagr |
| MTR 29592 | tararagr | tararagr | tararagr | tararagr |
| MTR 29566 | tararagr | tararagr | tararagr | tararagr |
| MTR 29586 | tararagr | tararagr | tararagr | tararagr |
| AMCC 204421 | tararagr | tararagr | tararagr | tararagr |
| MTR 29611 | tararagr | tararagr | tararagr | tararagr |
| MTR 29619 | tararagr | tararagr | tararagr | tararagr |
| MTR 29622 | tararagr | tararagr | tararagr | tararagr |
| MTR 29623 | tararagr | tararagr | tararagr | tararagr |
| CHUNB 74156 | tararagr | tararagr | tararagr | tararagr |
| MTR 29588 | tararagr | tararagr | tararagr | tararagr |
| CHUNB 74157 | tararagr | tararagr | tararagr | tararagr |
| CHUNB 74163 | tararagr | tararagr | tararagr | tararagr |
| AMCC 204364 | teyumirim | teyumirim | guarani + teyumirim | teyumirim |
| AMCC 204357 | guaranigr | guarani | guarani + teyumirim | guarani |
| AMCC 204453 | guaranigr | guarani | guarani + teyumirim | guarani |
| MTR 29373 | guaranigr | guarani | guarani + teyumirim | guarani |
| AMCC 204349 | guaranigr | guarani | guarani + teyumirim | guarani |
| MTR 29393 | guaranigr | guarani | guarani + teyumirim | guaira |
| MTR 29379 | guaranigr | guarani | guarani + teyumirim | acahay |
| AMCC 204510 | guaranigr | spinulosus | spinulosus | spinulosus |
| AMCC 204498 | guaranigr | spinulosus | spinulosus | spinulosus |
| AMCC 204481 | guaranigr | spinulosus | spinulosus | spinulosus |
| AMCC 204384 | guaranigr | spinulosus | spinulosus | spinulosus |
| LJAM-CNP 12113 | guaranigr | spinulosus | spinulosus | spinulosus |
| LG1551/2987 | xanthochilus | xanthochilus | xanthochilus | xanthochilus |
| AMCC 204537 | xanthochilus | xanthochilus | xanthochilus | xanthochilus |
| MTR 29503 | xanthochilus | xanthochilus | xanthochilus | xanthochilus |
| CHUNB 74137 | xanthochilus | xanthochilus | xanthochilus | xanthochilus |
| MTR 29466 | xanthochilus | xanthochilus | xanthochilus | xanthochilus |

| | | | | |
|---|---|---|---|---|
| MTR 29508 | xanthochilus | xanthochilus | xanthochilus | xanthochilus |
| CHUNB 74145 | xanthochilus | xanthochilus | xanthochilus | xanthochilus |
| MTR 29554 | xanthochilus | xanthochilus | xanthochilus | xanthochilus |
| MTR 29499 | xanthochilus | xanthochilus | xanthochilus | xanthochilus |
| MTR 29439 | xanthochilus | xanthochilus | xanthochilus | xanthochilus |
| 946104 | xanthochilus | xanthochilus | xanthochilus | xanthochilus |
| CHUNB 74146 | xanthochilus | xanthochilus | xanthochilus | xanthochilus |
| MTR 29521 | spnov | spnov | spnov | spnov |
| MTR 29440 | callathelys (Outgroup) | callathelys (Outgroup) | callathelys (Outgroup) | callathelys (Outgroup) |
